# Transposable element products, functions, and regulatory networks in Arabidopsis

**DOI:** 10.1101/2024.04.02.587720

**Authors:** Carles Borredá, Basile Leduque, Vincent Colot, Leandro Quadrana

**Affiliations:** Institute of Plant Sciences Paris-Saclay (IPS2), Université Paris-Saclay, CNRS, INRAE, Université Evry, Université Paris Diderot, 91190 Gif sur Yvette, France.; Institut de Biologie de l’Ecole Normale Supérieure (IBENS), Centre National de la Recherche Scientifique (CNRS), Institut National de la Santé et de la Recherche Médicale (INSERM), École Normale Supérieure, PSL Research University, Paris, France.

## Abstract

Transposable elements (TEs) are DNA sequences with the ability to propagate themselves within genomes. Their mobilization is catalyzed by self-encoded factors, yet these factors have been poorly investigated. Here, we leveraged extensive long-and short-read transcriptome data, structural predictions, and regulatory networks analyses, to construct a comprehensive atlas of TE transcripts and their encoded products in the model organism *Arabidopsis thaliana*. We uncovered hundreds of transcriptionally competent TEs, each potentially encoding multiple proteins either through distinct genes, alternative splicing, or post-translational processing. Structural-based protein analyses revealed hitherto unidentified domains, uncovering proteins with multimerization and DNA binding domains forming macromolecular complexes. Furthermore, we demonstrate that TE expression is highly intertwined with the transcriptional network of cellular genes, and identified transcription factors and cis-regulatory elements associated with their coordinated expression during development or in response to environmental cues. This comprehensive functional atlas provides a valuable resource for studying the mechanisms involved in transposition and their consequences for genome and organismal function.

## INTRODUCTION

Transposable elements (TEs) are ubiquitous DNA sequences that move and self-propagate across the genome. Although TEs pose a threat for genome integrity, they are also major drivers of evolution. For instance, through so-called “domestication”, TEs have enabled the emergence of proteins with important cellular functions, including the repurposing of TE transposases to give rise to the RAG recombinases in jawed vertebrates (Vladimir V. Kapitonov and Jurka 2005) or the transcription factors FHY3 and FAR1 that modulate light signaling in plants (Lin et al. 2007). In addition, cis-regulatory motifs within TEs have played important roles in the evolutionary rewiring of gene regulatory networks, such as in the case of the interferon response in mammals (Chuong, Elde, and Feschotte 2016). Despite the extensive knowledge regarding the contribution of TEs at the macroevolutionary scale to cellular functions and gene regulation, our understanding of factors encoded by the TEs themselves remains limited.

TEs are categorized in two broad classes based on their mobilization mechanisms. Class I retrotransposons move via an RNA intermediate, while Class II DNA transposons use a “cut and paste” mechanism instead. TEs are further divided into superfamilies and families based on particular sequence features, such as the presence of specific terminal repeats or conserved protein domains (Wicker et al. 2007). Transcription of TEs and subsequent synthesis of functional TE-encoded proteins are the first steps for their transposition. So-called non-autonomous TEs have at least one of these processes impaired, and rely for their mobilization on factors encoded by other TEs. Autonomous TEs often encode multiple proteins involved in transposition, either as separate transcription units, alternative splicing isoforms, or polyproteins, depending on the TE families considered. Investigating TE-encoded proteins has remained extremely challenging due to the dearth of reliable annotations of TE transcripts in reference genomes. Indeed, potentially mobile TE copies are typically kept transcriptionally silent through robust epigenetic mechanisms, hence the dramatical underrepresentation of *bona fide* TE gene and TE-encoded proteins in databases. One promising way to fill this gap is to make use of mutants deficient in factors involved in the epigenetic silencing of TEs, as was done in the model plant species *Arabidopsis thaliana* to generate a “gene-like” TE annotation (Panda and Slotkin 2020). However, due to the limited number and size of the full-length transcripts obtained so far, our current knowledge of the TE transcriptome is far from complete even in this species.

Here, by combining high quality long-and short-read transcriptomic data from *A. thaliana* plants defective in one or multiple layers of epigenetic silencing, we built a comprehensive atlas of TE-genes, determined the repertoire of TE-encoded protein products and functional domains, and explored the regulation of their expression. Structure-based analysis uncovered a large diversity of protein domains, many of which were not reported before. Moreover, we identified novel DNA binding and dimerization modules and we provide evidence for their interaction to form multiprotein complexes. We also show that TE expression is highly intertwined with the transcriptional network of genes, and we identified in numerous cases the transcription factors as well as the cis-regulatory sequences likely involved. Overall, our study provides a first comprehensive characterization of TE-genes, transcript isoforms, and proteins, as well as the mechanisms regulating their activity, thus greatly advancing our understanding of the functional significance of mobile TE sequences in extant genomes. This resource and the methodological framework presented pave the way for similar studies in other organisms.

## RESULTS

### High quality functional annotation of transposable element-encoded genes

To characterize the transcriptome diversity of TEs in *A. thaliana*, we obtained high quality ONT-based long-read cDNA sequencing data using poly(A)-containing RNAs extracted from plants defective in the chromatin remodeler DDM1, which is required for the DNA methylation-dependent silencing of TEs. In addition, we included plants combining the *ddm1* mutation with either a mutation in the gene encoding RNA dependent RNA polymerase 2 (RDR2) or RDR6, which are involved in siRNA-mediated transcriptional and post-transcriptional silencing of TEs, respectively (Figure 1A). We obtained more than 9 million reads, with a median read length of 1500 nucleotides (Figure S1 A). High-confidence transcript isoforms (Figure 1B) were identified using the Full-Length Alternative Isoform analysis of RNA (FLAIR) bioinformatic pipeline (Tang et al. 2020). In total, we assembled 40,710 transcript isoforms, which overlap 16,735 cellular genes and 1,038 TEs annotated in the TAIR10 reference genome (Figure 1C). 99.6% of exon-exon junctions identified by our approach over cellular genes were validated by Illumina short-reads obtained from 462 RNAseq experiments (supplementary table 1)(Figure S1 B). Furthermore, almost all exon boundaries (98.61%) and isoforms (87.11%) coincided with those annotated in the TAIR10 reference genome or the splicing-aware annotation AtRTD2 (Zhang et al. 2017) (Figure S1 C and D). In addition, our predicted transcript orientation matched the TAIR10 annotation in 97.4% of the cellular genes (Figure S2 A), and the stranded Illumina transcriptomes validated 97.9% of them (Figure S2 B). Altogether, these results demonstrate the high specificity of our *de novo* transcript annotation.

**Figure 1.**
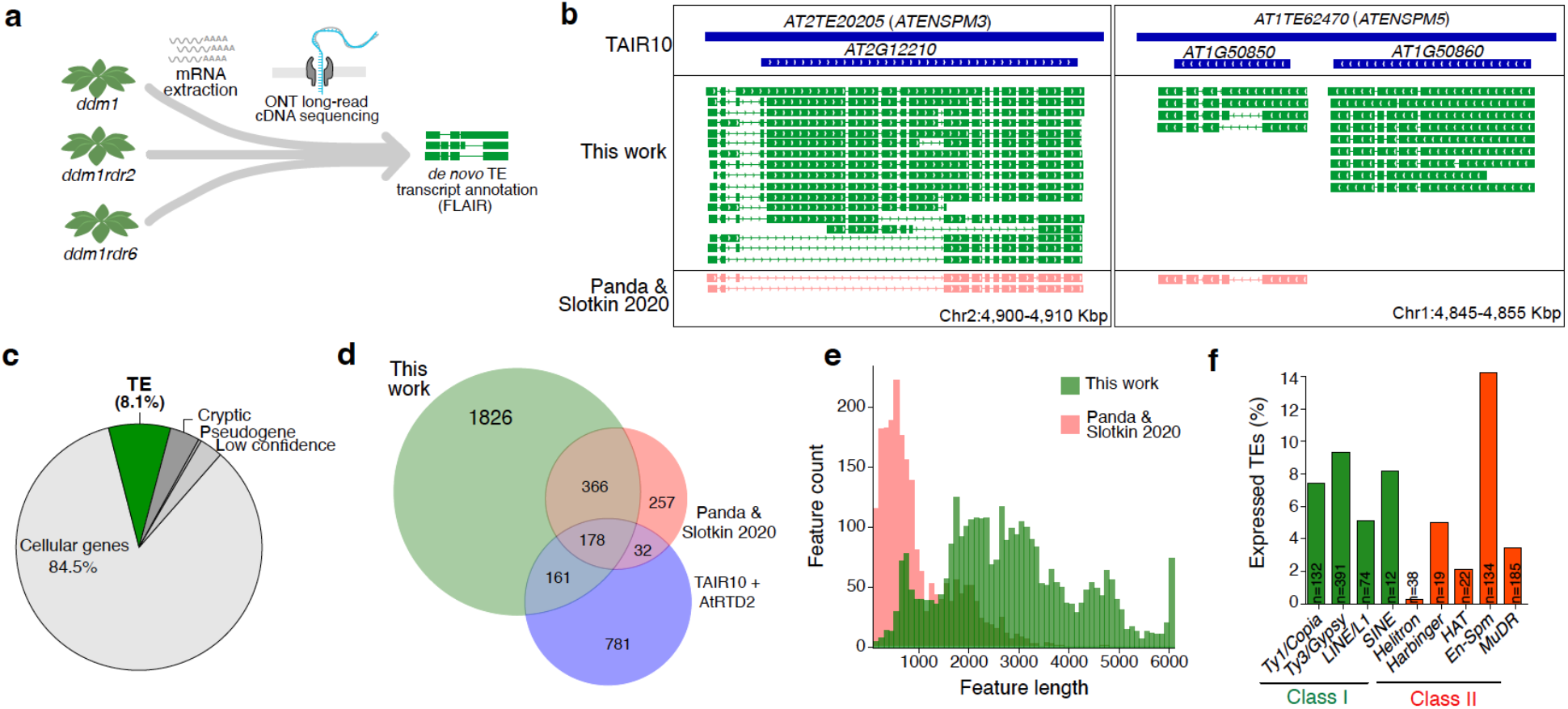
A high-quality annotation of transposable elements-encoded genes. **a**. Overview of the experimental workflow and annotation pipeline used. **b**. Selected examples of transposable element gene annotations from previous annotation efforts compared to our study. **c**. Proportion of FLAIR-annotated transcripts classified according to their genomic origin. **d**. Shared and private TE-gene intron-containing isoforms annotated in this work compared with previous annotation efforts. Isoforms are considered shared when they contain the exact same set of exon-exon junctions. **e**. Length distribution of TE-gene isoforms annotated in this work compared with previous studies. **f**. Proportion of transcriptionally competent TEs per superfamily. The total number of competent TEs is displayed.

We next characterized the transcripts generated exclusively by TEs, as opposed to chimeric transcripts that span both TEs and cellular genes (Berthelier et al. 2023). In total we identified 3,410 TE-derived transcripts, 76.8% of which contain at least two exons. The majority of transcripts derive from copies belonging to the LTR retrotransposon superfamily *Gypsy/Ty3*, followed by the DNA transposon superfamilies *MuDR* and *SPM,* as well as the LTR retrotransposon superfamily *Copia/Ty1* (Figure 1F), consistent with many of the corresponding families being relatively young (V. V. Kapitonov and Jurka 1999). Importantly, most of the identified TE transcripts were not annotated in the reference TAIR10 genome nor reported before (Panda and Slotkin 2020), likely due to the difficulties to detect long and/or lowly expressed transcripts with short-or early long-read sequencing methods (Figure 1E and Figure S3). Notably, only 13.4% of all TE transcript isoforms and 34% of all exon-exon junctions detected by our long-reads matched previous TAIR10 or AtRTD2 annotations (Figure 1D), suggesting that most TE-encoded transcripts are missing or mistaken in currently available annotations. Orientation of our long-read based TE transcripts matched TAIR10 TE-gene annotation in 77% of the cases (Figure S2 A). For the remaining 33%, stranded RNA-seq data supported our prediction in the majority of cases (Figure S2 B). Overall, we estimate that 80% of TE-derived transcripts that we have detected are incorrectly annotated or altogether missing in the TAIR10 reference genome.

### The transposable elements transcriptome

Transposition typically requires the activity of multiple protein factors, which may be encoded in different genes within a TE copy (Wicker et al. 2003; Hershberger et al. 1995), or else produced by alternative splicing (Masson et al. 1989) or post-translational processing of immature proteins (Youngren et al. 1988). Analysis of our transcripts revealed that the number of non-overlapping transcription units harbored by TEs (hereafter referred as TE genes) varies greatly among, as well as within, TE superfamilies. For instance, *MuDR* DNA transposons generally contain two to three TE genes, with some loci containing up to six (Figure 2A). Conversely, *SPM3* and *SPM7* DNA transposons contain a single TE gene (Figure 2B and Figure S4), reminiscent of the maize DNA transposon *Spm*, which contains a single gene that generates two alternative transcript isoforms encoding either the TnpD transposase or else the potential regulatory factor TnpA (Masson et al. 1989). Unexpectedly, we found several Arabidopsis *SPM* families (*SPM2, 5, 6* and *9*) containing two non-overlapping head-to-tail genes (Figure S4), likely encoding putative TnpD and TnpA factors (Figure S5). In addition, we identified three families, *SPM1*, *1A,* and *4,* which encode for TnpAs closely related to the one encoded by a full-length *SPM2*. The absence of TnpD and similarity between TnpAs among these families suggest that *SPM1*, *1A,* and *4* are non-autonomous and rely on SPM2-encoded TnpA for their propagation. Altogether, these results illustrate the large diversity of gene architectures even among closely related TE families.

**Figure 2:**
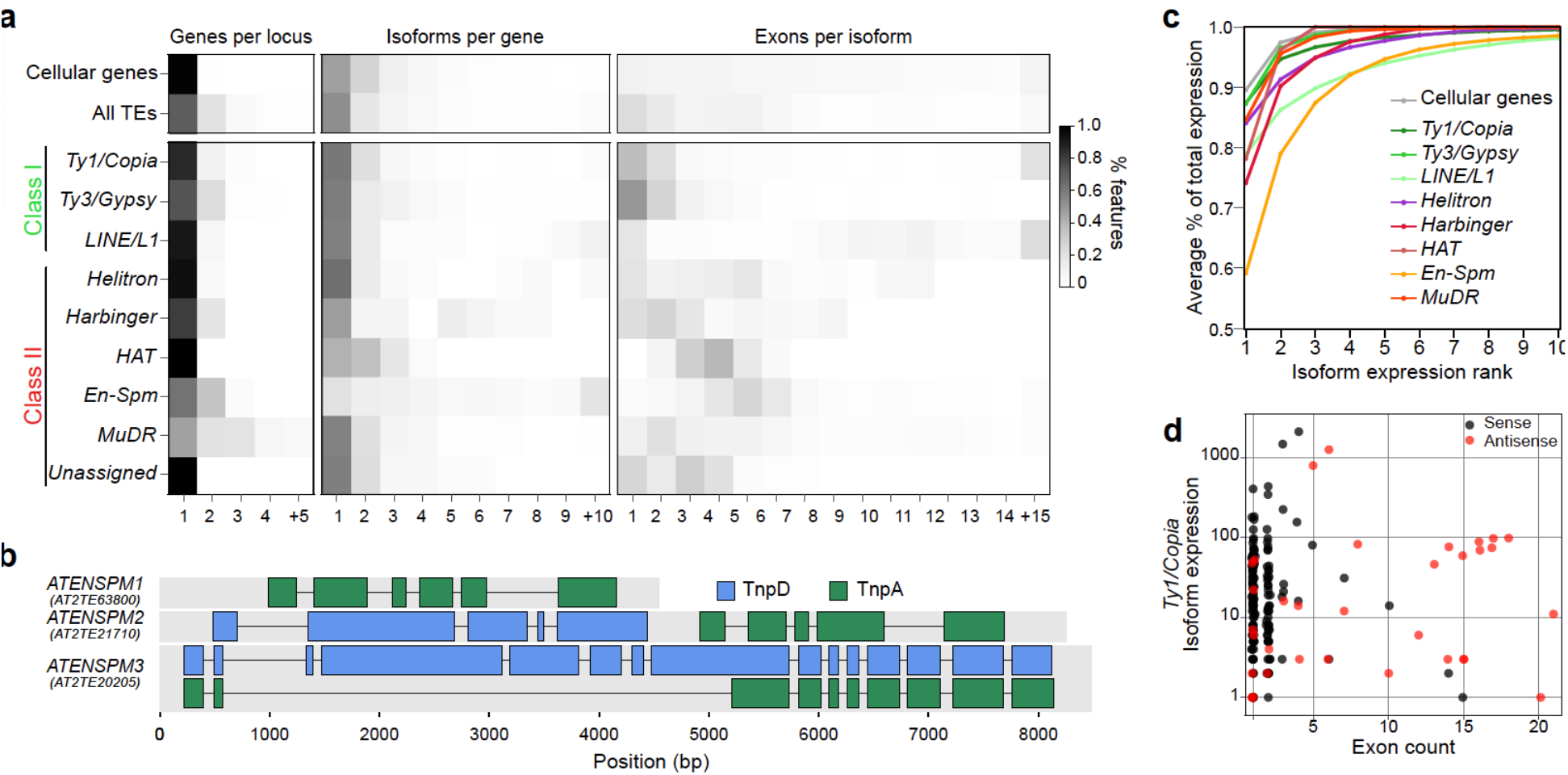
The transposable elements transcriptome. **a.** Relative distribution of annotation features from cellular and TE-genes (top) and for different TE superfamily. **b**. Protein-coding isoforms of selected SPM families. TnpA and TnpD encoding genes and isoforms are shown in green and blue, respectively. Only the isoform coding the longest ORF for TnpA and TnpD is displayed. **c**. Average contribution to total expression per ranked isoform for each TE superfamily. **d**. Absolute expression of sense and antisense *Ty1/Copia* transcripts.

TE genes generate fewer transcript isoforms compared to cellular genes, with the notable exception of *SPM* and *LINE* elements. For example, *AT3G32230* (*SPM2)* and *AT2G10250* (*ATLINE1_6)* TE-genes are capable of producing over 37 distinct transcript isoforms each. Nonetheless, one or two isoforms typically account for the majority of steady state mRNAs (Figure 2C).

*Copia/Ty1* and *Gypsy/Ty3* usually contain only one gene producing a long transcript that covers most of the TE length, consistent with these full-length mRNAs serving as templates for retrotranscription and encoding the polyprotein precursors (Youngren et al. 1988). We also identified 51 *Copia/Ty1* retrotransposon copies that produce alternative spliced transcript isoforms, many of them encoding the GAG protein as previously reported in Arabidopsis (Oberlin et al. 2017). More intriguingly, we noted numerous multi-exonic transcripts associated with a subset of phylogenetically related *Copia/Ty1* LTR-retrotransposons (Figure S6). These multi-exonic transcripts were antisense, non-coding, and generally initiated within the 3’ LTR (Figure S7). Expression levels of antisense and sense transcripts were comparable (Figure 2D), suggesting that the former are not produced by spurious transcription. In fact, the conservation of these antisense transcripts across several related TE families suggest that they may play a functional, potentially regulatory, role. Taken together, our findings underscore the diverse mechanisms TEs employ to encode the multiple functions crucial for their propagation.

### A hidden diversity of TE-encoded structural domains

Next, we sought to determine the coding potential of the 3211 high-quality TE-encoded transcripts we identified. 2953 of these transcripts (91.9%) contain ORFs encoding peptides longer than 50 amino acids. About one third of these predicted peptides contain at least one conserved domain (CD) in the CDD database (Figure S8). Of note, we did not find any HELITRON with relevant ORFs, thus suggesting that none of these TEs are mobile or else leaving open the question of the factors involved in the mobilization of members of this intriguing superfamily of TEs (Barro-Trastoy and Köhler 2024).

The fast evolving rate of TE-encoding proteins compared to cellular proteins limits the ability of sequence-based CD-searches to identify conserved domains. Protein structures instead retain homology for much longer than primary sequences, facilitating the identification of conserved domains (Illergård, Ardell, and Elofsson 2009). Hence, structural-based analysis holds great promise to investigate the functions and evolution of TE-encoded products. However, TE-encoded proteins are systematically underrepresented in protein structure databases, including the recently established AlphaFold Protein Structure Database, which comprises over 200 million protein folding predictions (Varadi et al. 2022). We therefore set out to predict the structure of the 2953 high-quality TE-encoded putative proteins using AphaFold2 (Jumper et al. 2021). High or very high confidence (i.e. pLDDT>70) structural predictions were obtained for most TE-encoded protein sequences (Figure S9). In total, we identified 3131 compact structural domains (SD) across the 2953 high-quality putative TE-encoded proteins. Clustering of these SDs based on structure similarity yielded 201 clusters containing at least two proteins, and 737 singleton SDs that likely represent chimeric or orphan peptides (Figure S10). Clustering of these SDs allows us to reconstruct representative structure and domain architecture of all TE-encoded proteins in Arabidopsis (Figure 3).

**Figure 3:**
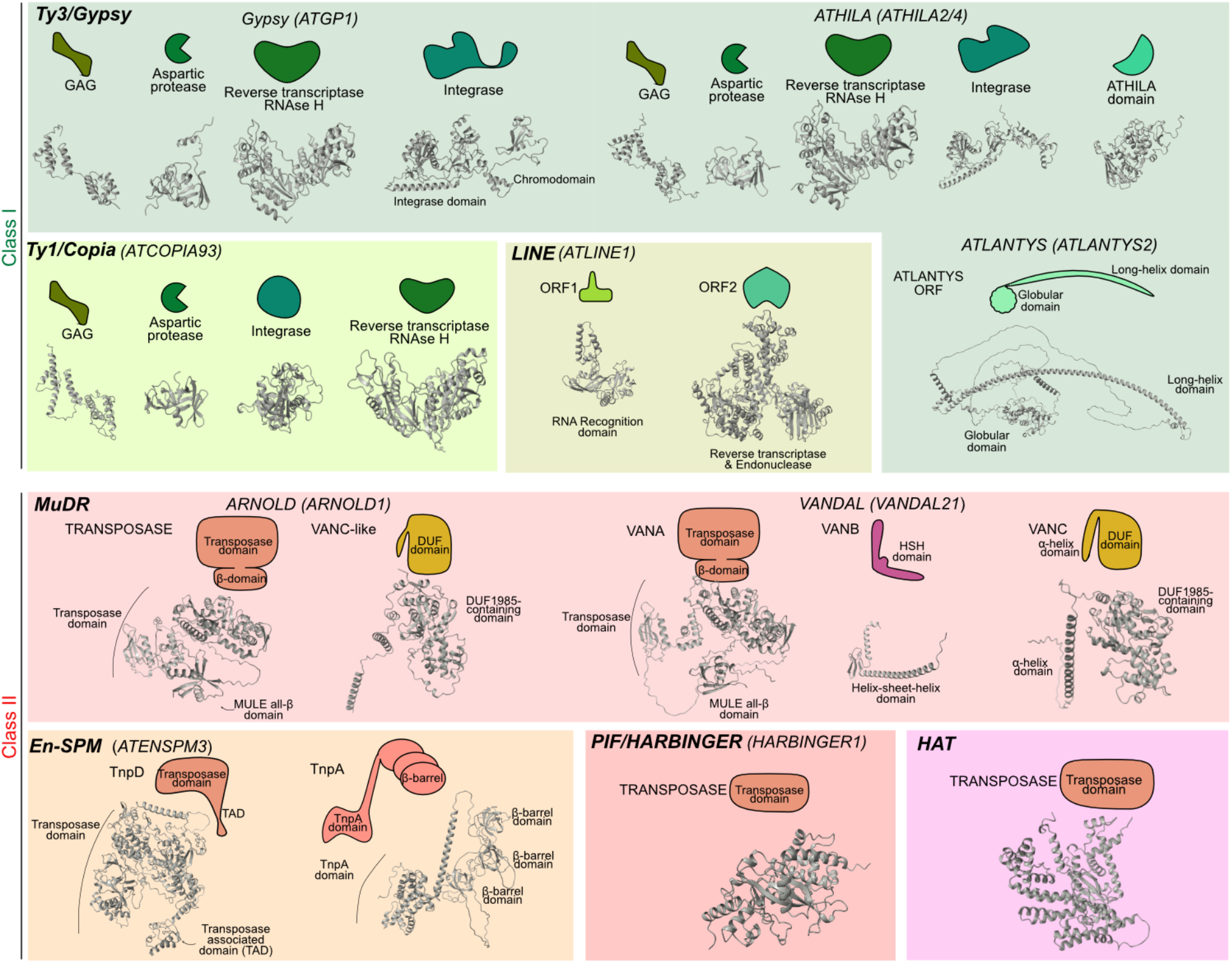
Transposon-encoded protein architectures. Representative proteins, domains, and architectures are indicated for the different major transposon superfamilies.

We next sought to characterize the putative function and origin of the 938 different types of TE-encoded SDs (Figure 4A). To this end, we compared their structure with all experimentally solved structures available at the Research Collaboratory for Structural Bioinformatics PDB (RCSB PDB) database, as well as the predicted structures in the AlphaFold Protein Structure databases, using FoldSeek (van Kempen et al. 2023). In total, we annotated 721 SDs, comprising 83.3% of all putative TE-encoded proteins we identified. A large fraction of these SDs have significant structural hits with cellular proteins. For example, *ATLANTYS* LTR retrotransposons encode a SD with strong structural homology with the meiotic–specific Asynaptic 2 (ASY2) protein (Figure 4B) implicated in the formation of meiotic chromosome axis via coiled-coil oligomerization (West et al. 2019; Armstrong et al. 2002). We also identified an SD within the SPM-encoded TnpD transposase that has structural homology to the DNA binding domain of the mouse RAG recombinase. The RAG recombinase, derived from a *Transib* transposon, cleaves DNA through a nick–hairpin mechanism, stimulating non-homologous DNA end joining (Lieber et al. 2004; Huang et al. 2016). The significant structural parallels between RAG1 and TnpD suggest similar cleavage mechanisms, offering a plausible explanation to the high rate of excision observed experimentally for *SPM3* (Quadrana et al. 2019). In addition, we found that the C-terminal region of *SPM3*-encoded TnpA contains a string of three globular SDs (Figure 4D). These SDs folds are composed of five short antiparallel beta sheets forming a small β-barrel, a ubiquitous folding associated with nucleic acid binding and chromatin organization (Youkharibache et al. 2019). Moreover, the structure of TnpA’s β-barrel domains resemble the NDX-B domain of the plant-specific transcription factor NODULIN HOMEOBOX PROTEIN (Mukherjee, Brocchieri, and Bürglin 2009), which interacts with core components of the Polycomb Repressive Complex 1 (Zhu et al. 2020). This similarity raises the intriguing possibility that TnpA factors regulate *SPM* transposition via an interaction with PRC1.

**Figure 4:**
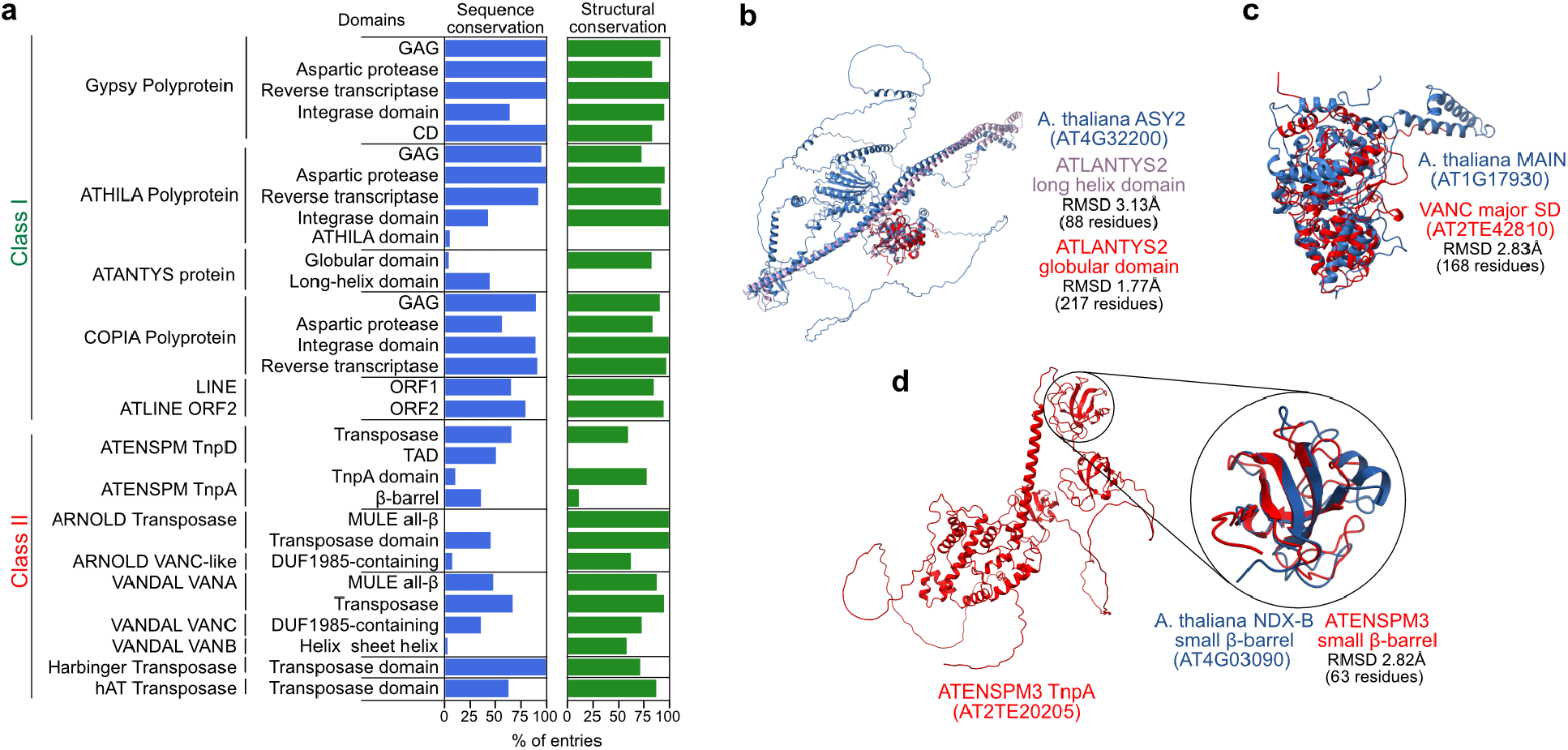
Structure-driven characterization of transposon proteins. **a.** Proportion of TE-encoded structural domains with hits in conserved domains or structural similarity to proteins from public databases. **b**. Structural alignment of ATLANTYS-encoded globular and long-helix domains (red and purple, respectively) with MEIOTIC SYNAPTIC MUTANT 2 ASY2 protein (blue). **c.** Structural alignment of the DUF-containing domain of VANC (red) and MAINTENANCE OF MERISTEMS (MAIN) protein (blue). **d**. Predicted structure of *ATENSPM3*-encoded TnpA protein (in red). Structural alignment of the first β-barrel and the NDX-B β-barrel (in blue) is displayed.

TE-encoded chromatin regulation has also been described for *VANDAL* DNA transposons, which encode the anti-silencing factors VANC (Sasaki et al. 2022; Fu et al. 2013; Hosaka et al. 2017). Our structural homology search revealed strong similarity between the largest SD of VANDAL-encoded VANC, ARNOLD-encoded unknown proteins, and the cellular factor MAINTENANCE OF MERISTEMS-like protein 1 (MAIL1) (Figure 4C), which is involved in chromatin condensation and TE silencing (Ikeda et al. 2017). Since the emergence of MAIL1 predates that of VANC (Sasaki et al. 2022; Ikeda et al. 2017), *VANDAL* and ARNOLD elements, which belong to a superfamily of TEs (*MuDR*) that is prone to capture cellular genes (Jiang et al. 2004), might have acquired MAIN1-like domains and repurposed them into anti-silencing factors.

Arabidopsis heterochromatin is densely populated by *Ty3/Gypsy* LTR-retrotransposons belonging to five different *ATHILA* families, which have been recently proposed to play a role in centromere evolution (Wlodzimierz et al. 2023). The reference Arabidopsis genome (Col-0) harbors many centromeric as well as non-centromeric *ATHILA* copies, all of which are thought to be inactive. Supporting this notion, we found that none of 145 the *ATHILA* elements identified in our dataset encode full-length polyproteins carrying the canonical structural domains: GAG protein, aspartic protease, retrotranscriptase/RNAseH, and integrase. In contrast, we identified at least 13 *Ty1/Copia* and three non-*ATHILA Ty3/Gypsy* elements encoding full-length polyproteins harboring the complete set of retrotransposition factors. Coding of multiple proteins from the same transcript can also be achieved by ribosomal frameshift, as is the case for several retroviruses and yeast retrotransposons (Belcourt and Farabaugh 1990). We thus tested for the presence of multiple ORFs within LTR-retroelements that may be indicative of ribosomal slippage. We found a single *ATGP3* element (*AT1TE42395*) encoding a 0-frame full-length GAG protein and a-1-frame polyprotein harboring the aspartic protease, retrotranscriptase/RNAseH, and integrase SDs. In combination, these results support the absence of functional *ATHILA* copies in the Arabidopsis reference genome and provide the first evidence for potential ribosomal frameshifting of *Ty3/Gypsy* retrotransposons in Arabidopsis (Figure S11).

Overall, we identified at least 67 TEs belonging to 34 families, belonging to *Ty1/Copia*, *Ty3/Gypsy*, *MuDR*, *SPM*, *hAT*, *Harbinger* and *LINE/L1* superfamilies, producing the complete set of full-length proteins required for transposition (Supplementary table 2). This number is likely an underestimation, as it only encompasses TEs expressed in the tissues and conditions tested.

### Disordered C-terminal domains of integrases carry diverse targeting signals

Besides SDs, our structural predictions yielded a large number of low confidence foldings (Figure S9), indicative of intrinsically disordered regions. In particular, we noted that the single protein encoded by *Ty1/Copia* and *Ty3/Gypsy* LTR retrotransposons are typically composed of four or five large SDs linked by intrinsically disordered regions. Processing of immature polyprotein by the self-encoded aspartic protease releases the different mobility factors, including the integrase, GAG, and retrotranscriptase(Youngren et al. 1988). In yeast, retrotransposon-encoded aspartic proteases target patterns of hydrophilic/hydrophobic residues (Merkulov et al. 1996; Kirchner and Sandmeyer 1993), however the ubiquity of such degenerated patterns hinders the prediction of potential cleavage sites. We leveraged the structural information of SD boundaries together with hydrophilic/hydrophobic pattern searches to predict protease cleavage sites (see methods) across 74 retrotransposon-encoded polyproteins containing integrases. We identified 29 *Ty1/Copia* and 13 *Ty3/Gypsy* (Figure S12) full-length integrases harboring intrinsically disordered and highly divergent C-terminal domains (CTD). CTDs of retroviral as well as yeast *Ty1* integrase were shown to be essential for nuclear localization (Moore, Rinckel, and Garfinkel 1998) and integration targeting (Asif-Laidin et al. 2020). We thus tested for the presence of nuclear localization signals (NLS) across the 29 *Ty1/Copia* and 13 *Ty3/Gypsy* CTDs. Despite the extensive sequence divergence, we found that two thirds of *Ty1/Copia* integrases contain monopartite NLS motifs, while most *Ty3/Gypsy* elements carry bipartite NLS (Figure S12). Variation in the type of NLS signals among retrotransposon classes may affect their interactions with importing proteins, potentially modulating their nuclear sorting and localization. The identification of protease cleavage sites and CTDs of integrases thus opens the possibility to investigate the regulation, subcellular sorting, and insertion preferences of integrases encoded by LTR-retrotransposons.

### TE-encoded factors typically interact to form multiprotein complexes

TE mobilization typically involves multiprotein complexes, however the extent of TE-encoded protein-protein interaction is unknown. We next investigated the potential ability of TE-encoded proteins to produce homo-and heterodimers using AlphaFold-Multimer (Evans et al. 2022). We focused on 17 proteins encoded by a representative subset of potentially autonomous TEs (Supplementary table 2) spanning the different superfamilies and for which mobilization has been observed experimentally (Quadrana et al. 2019) or inferred in nature (Baduel et al. 2021). In contrast to the overall scarcity of predicted heterodimers (8/136 possibilities), two thirds (11/17) of the TE-proteins analyzed produce homodimers, with the exception of the transposase VANA (*VANDAL*), Retrotranscriptase/RNAseH (*Copia/Ty1* and *Gypsy/Ty3*), ORF1 and ORF2 (*LINE*), and *Copia/Ty1* Integrases (Figure 5A). Further characterization of homodimerization domains revealed that the integrase of the *Ty3/Gypsy* retrotransposon *ATGP3* (*ATGP3:AT3TE16035*) has a large interacting surface involving scattered residues (Figure 5B), consistent with previous observations (Abascal-Palacios et al. 2021). Similarly, the protease encoded by *Ty1/Copia ATCOPIA93* (*ATCOPIA93:AT5TE20395*) displayed a large interaction surface formed by scattered residues (Figure 5C). Conversely, predicted homodimerization of the anti-silencing factor VANC21 (*VANDAL21:AT2TE42810*) involves a narrow protein region spanning an uncharacterized coiled-coil alpha-helix domain (Figure 5D). Likewise, predicted dimerization of the putative regulatory TnpA-like protein of *SPM3* also relies on a 40 amino-acid-long coiled-coil structure (Figure 5E and Figure S13). Given the independent origin of VANC and TnpA proteins, our results suggest that homodimerization through coiled-coil domains evolved convergently.

**Figure 5:**
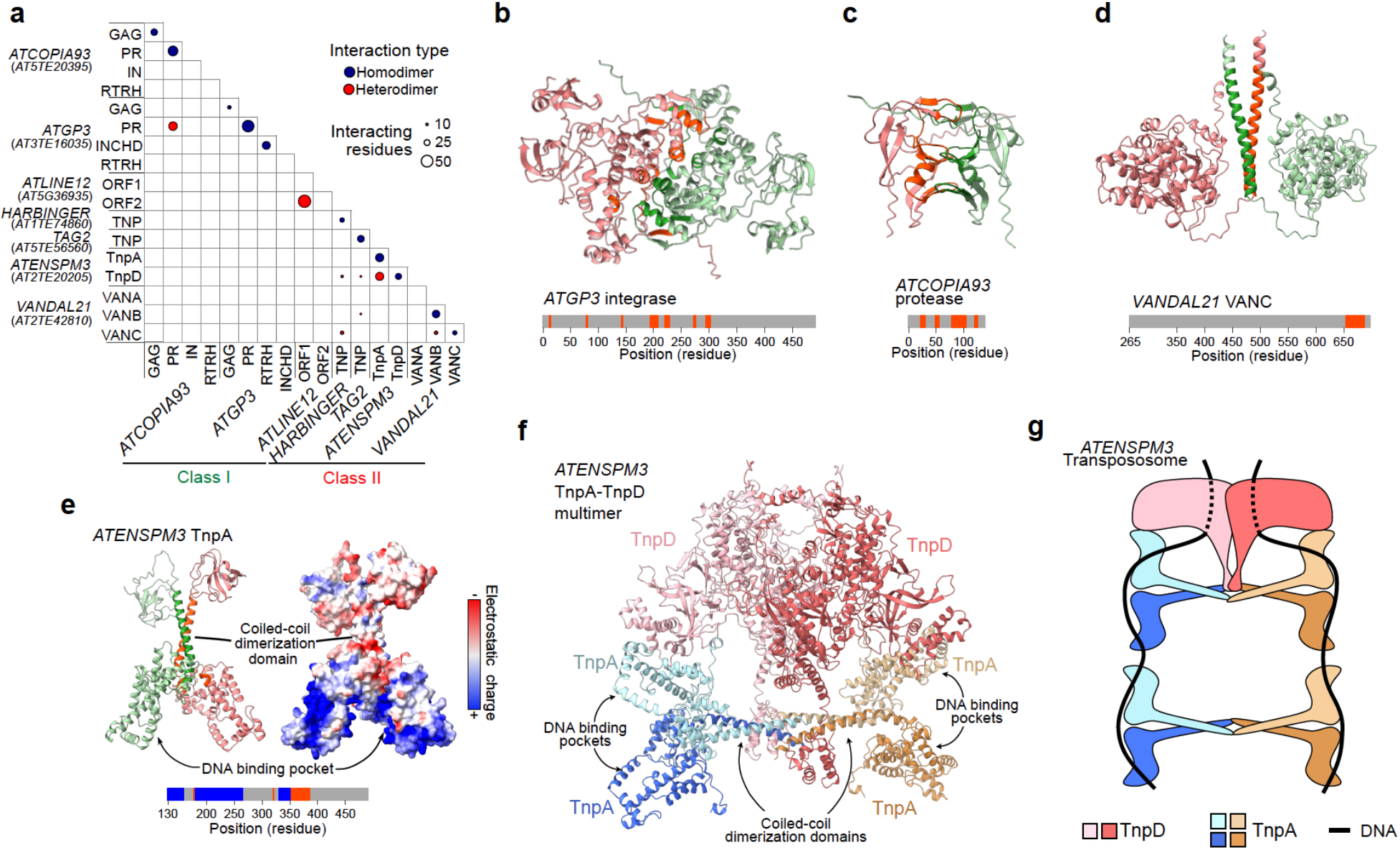
Interaction landscape of transposon proteins. **a**. Interaction matrix among a representative set of TE-encoded proteins. Blue and red dots represent homo and heterodimerization events, respectively. Dot sizes are proportional to the number of interacting residues. **b**. Predicted dimer conformation of *ATGP3*-encoded integrase. **c**. Predicted dimer conformation of *ATCOPIA93*-encoded protease. **d**. Predicted dimer conformation of *VANDAL21*-encoded VANC protein. The disordered N-terminal tail was hidden for clarity. **e**. Predicted dimer conformation of *ATENSPM3*-encoded TnpA protein and its electrostatic surface. Residues mediating the dimerization or located in the predicted DNA binding pocket are shown in orange and blue, respectively. The disordered N-terminal tail and the β-barrel domains are hidden. **f**. Predicted heteromultimer structure of *ATENSPM3*-encoded TnpA and TnpD proteins. TnpA disordered N-terminal region and β-barrel domains are hidden. **g**. Schematic representation of *ATENSPM3* transpososome model.

In addition to homodimerization, *SPM3* TnpA-like protein shows strong heterodimerization potential with its cognate transposase TnpD. Based on biochemical observations, interactions between maize TnpA and TnpD have been proposed to form macromolecular complexes (Raina et al. 1998). Indeed, we predicted that TnpD dimers additionally interact with several TnpAs to form a multiprotein complex (Figure 5F). The largest SD of TnpA contains a positively-charged pocket indicative of DNA binding (Figure S13), which we further confirmed by structural-based predictions (Yuan et al. 2022). In combination, the multimer structure and domain annotations suggest that TnpAs interact with the DNA and dimerize through their extended coiled-coil domains, potentially bridging TnpA-DNA complexes on different DNA molecules. This so-called synaptosome complex can additionally interact with TnpD transposases to form the multi-subunit protein-DNA transpososome (Figure 5G).

### Intertwined regulatory networks of TEs and genes

TEs are thought to have a remarkable ability to perceive developmental and environmental cues to achieve transposition (Fedoroff 1989). Nonetheless, comprehensive information regarding the transcriptional regulation of most TEs in response to endogenous or exogenous cues remains elusive. We exploited our high-quality TE-gene annotation to investigate 462 publicly available RNAseq data sets from wild-type Arabidopsis plants covering a wide range of tissues, developmental stages, and growth conditions (Supplementary table 1). Expression profile varies largely between TE families, providing opportunities to explore their transcriptional regulation (Figure S14). To systematically investigate the specificities of TE expression, we reconstructed the transcriptional network of 4,037 TE-and 28,445 cellular genes using co-expression clustering methods. Respectively 72% and 84% of TE-and cellular genes were grouped in 30 coexpressed clusters. Based on Gene Ontology enrichments, we assigned a representative biological function to each cluster (Figure S15). Intriguingly, TE-genes are spread across most clusters, spanning a wide diversity of biological functions (Figure 6A). The majority (64.2%) of coexpressed TEs and genes are non-adjacent (>1Kbp), which indicates that co-regulation due to genomic proximity (i.e. positional effects) is uncommon, and suggest instead that transcriptionally active TEs often contain regulatory elements similar or identical to that of genes (see next section). Thus, while it was previously reported based on fragmented transcriptomic data that TEs undergo global reactivation in pollen (Slotkin et al. 2009), our analysis revealed that only a small fraction of transcriptionally competent TEs are in fact active in male reproductive tissues (Figure 6B). Similarly, only few TEs appear to be upregulated in response to any given biotic or abiotic stress (Figure 6A and Figure S14). For instance, six full-length copies of the LTR-retrotransposon *ATCOPIA78/ONSEN* family, which represents a classical example of a stress-responsive retrotransposon (Ito et al. 2011), clustered together with 9 full-length TEs from different families and genes involved in abiotic stress response, and become reactivated in response to heat shock (Figure 6C). Similarly, we found several copies of the LTR-retrotransposon family *ATGP2N* within the defense response expression cluster (Figure 6D), indicating that this family becomes reactivated during pathogen infections. In summary, TE-genes display idiosyncratic responses to developmental and environmental cues.

**Figure 6.**
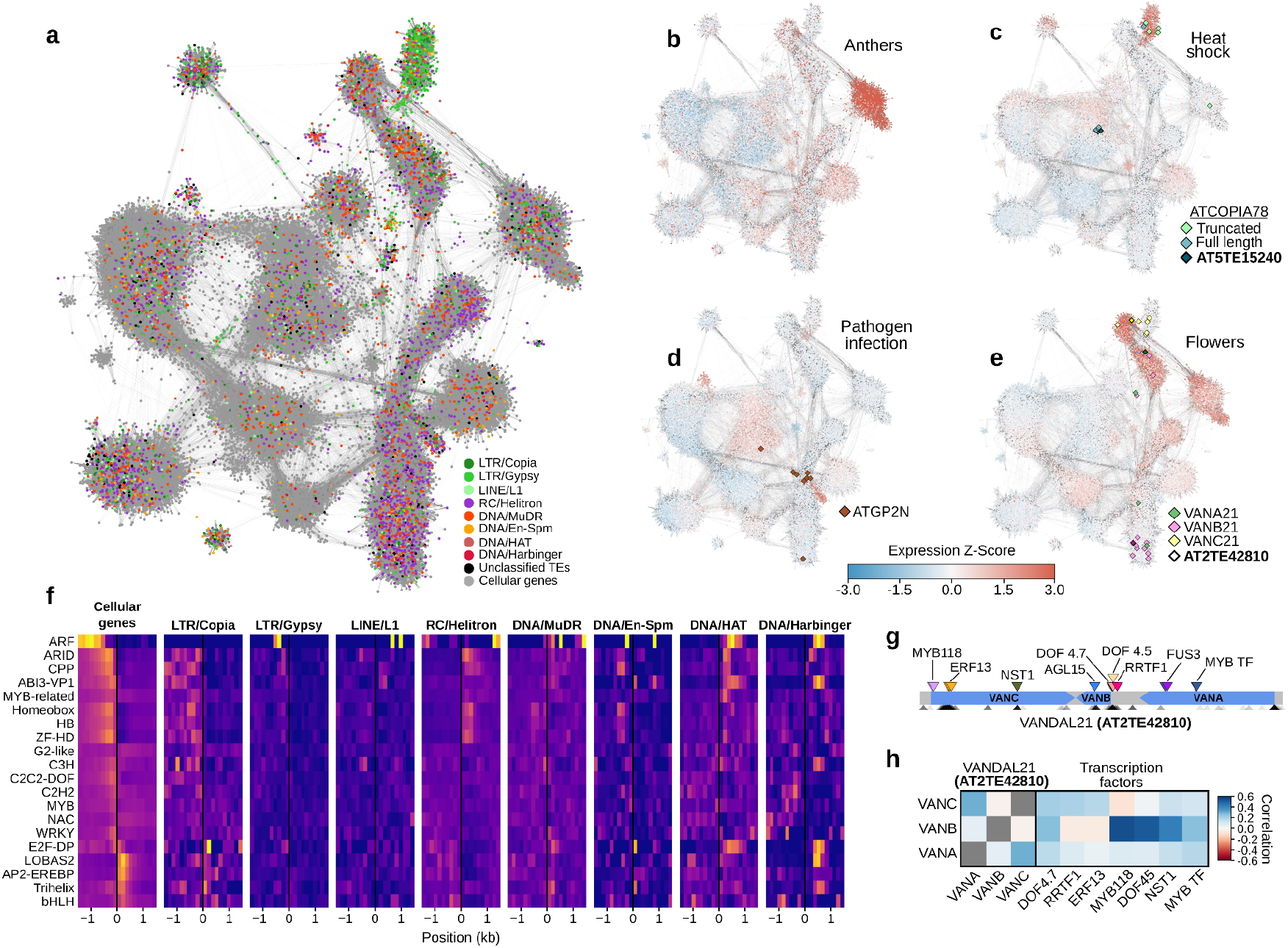
TE-gene expression networks. **a**. Coexpression network of TE-and cellular-genes. TE superfamilies and cellular genes are indicated with colors. **b**. Normalized expression of all cellular-and TE-genes in anthers. **c**. Normalized expression of all cellular-and TE-genes from leaves tissues subjected to heat shock. Full-length and truncated *ATCOPIA78* copies are shown. The mobile copy *ATCOPIA78 copy AT5TE15240* is highlighted. **d**. Normalized expression of all cellular-and TE-cellular genes in leaves after pathogen exposure. *ATGP2N* elements are highlighted. **e**. Normalized expression of all cellular-and TE-genes from flower tissues in control conditions. *VANA21*, *VANB21* and *VANC21* genes are shown. Genes encoded by the mobile VANDAL21 copy *Hiun* (*AT2TE42810*) are highlighted in bold. **f**. TFBS density of selected TF families around the TSS of cellular-andTE-genes. **g**. Distribution of TFBS along *Hiun* (*AT2TE42810*). TFBS for coexpressed and non-coexpressed TFs are presented on the top and the bottom, respectively. **h**. Expression correlation of *VANA21*, *VANB21* and *VANC21 genes* encoded by *Hiun* (*AT2TE42810*) and coregulated transcription factors with cognate TFBS. The three TFs with the strongest expression correlations for each TE-gene are shown.

We next set out to explore the transcriptional regulation of TEs carrying multiple genes. We focused on *VANDALs*, as TEs belonging to the *SPM* superfamily are barely expressed in any condition. We found that *VANA* and *VANC* genes of the *VANDAL21* copy *Hiun (AT2TE42810*) are both preferentially expressed in flowers (Figure 6E), suggesting concerted transcriptional regulation. Conversely, *Hiun*-encoded *VANB* gene, the function of which remains enigmatic, is located far away in the transcriptional network and coexpressed instead with genes involved in embryogenesis and germination. These results indicate that multigenic DNA transposons are subjected to complex transcriptional regulations leading to gene-specific expression patterns. To investigate the potential role of cis-regulatory elements in the regulation of TE-genes, we identified transcription factor binding sites (TFBS) within TE sequences by implementing a TE-aware analysis of *in vitro* DNA affinity purification sequencing (DAP-seq) data available for 529 Arabidopsis transcription factors (O’Malley et al. 2016). In total, we identified ∼4.5M putative TFBS within DAP-seq peaks, the majority of which were within or nearby genes and previously reported (Figure S16). However, almost half of the peaks identified exclusively by our method spanned TEs, offering a wider landscape of cis-regulatory elements over TES.. We found that retrotransposons, particularly *Ty1/Copia* and *Ty3/Gypsy*, tend to carry TFBS upstream of TSSs, consistent with LTR sequences potentially playing regulatory functions (Galindo-González et al. 2017). In contrast, TFBS at DNA transposons are often located downstream of TSS (Figure 6F), suggesting that transcriptional regulation of Class II TEs may rely on internal regulatory sequences.

To identify TFs directly involved in the regulation of TEs, we compared the expression profile of TFs with that of TE-genes carrying cognate TFBS. In total, we identified 1204 pairs of coexpressed TFs-TE-genes. TFBS for co-expressed TFs-TE were enriched closer to the TSS of co-regulated TEs compared to non-coexpressed pairs, particularly for Helitron and MuDR superfamilies (Figure S17). These coexpressed TFs-TE-genes provide the most relevant set of TFs likely regulating TE activity in Arabidopsis. For instance, we identified Dof-type zinc finger TFs associated exclusively with *VANDAL21*-encoded gene *VANB21*, while flower-specific *VANC21* and *VANA21* associate with Ethylene Response Factor 13 (Figure 6G and 6H), likely driving the tissue-specific expression regulation of these genes. In combination, our results demonstrate that different TE families, as well as genes within TE copies, evolved idiosyncratic transcriptional regulations through the presence of specific cis-regulatory sequences.

## DISCUSSION

Despite being major components of most eukaryotic genomes, TEs and their encoded products are overwhelmingly understudied compared to cellular genes. Our comprehensive analysis of TE transcripts and proteins provides a first integrated functional characterization of the large and diverse repertoire of active TEs in the *A. thaliana* genome. Given that many of these TEs families are widely shared across eukaryotes, our findings and methodologies are highly relevant to study TEs across the tree of life.

TE mobilization necessitates the synchronized activity of multiple TE-encoded factors. Our findings unveil the various mechanisms that distinct TEs exploit to encode multiple functions, such as carrying multiple genes, through alternative splicing, or post-translational processing. Strikingly, we uncovered that closely related TE families have switched between these mechanisms, suggesting that they can evolve remarkably fast. Furthermore, we demonstrate that polygenic *VANDAL* elements have complex transcriptional regulations, leading to contrasted tissue-specific expression patterns among different genes of the same TE copy. We further show that such idiosyncratic gene regulations are associated with specific cis-regulatory sequences within the TE themselves.

Our protein structure predictions for 2953 putative TE-encoded proteins revealed hundreds of TE-encoded structural domains that are absent from existing protein databases. This finding has broad implications for the study of TE evolution and their role in the emergence of new cellular functions. For instance, the excess of domesticated transposases found in vertebrates may reflect biases in sequence homology-based approaches, which are limited to detecting domains already annotated in databases. In fact, transposases are extremely ancient and their genes are among the most abundant in genomes (Aziz, Breitbart, and Edwards 2010). The expanded repertoire of TE-derived domains presented here may serve to clarify the evolution of seemingly orphan proteins, whose origin remains elusive. Thus, our findings pave the way for a better understanding of TE evolution and provide valuable insights into the origins and diversification of cellular functions.

By employing structural-guided analysis, we uncovered dozens of unknown structural domains, particularly within fast evolving proteins such as sequence-specific anti-silencing factors. Furthermore, we assigned functions to many of these unknown domains, including DNA binding pockets and multimerization domains in multiple retrotransposons-and DNA transposons-encoded proteins. Importantly, we observed much higher frequency of TE-encoded proteins forming homo-dimers (64%) compared to cellular genes, where less than 13% of proteins were found to form homodimers (Material et al. 2011). The high frequency of homodimers could reflect the unique capacity of TEs to encode, within remarkably constrained DNA sequence lengths, self-sufficient transposition machinery. (Moore, Rinckel, and Garfinkel 1998). A case in point are the aspartic proteases encoded by LTR-retrotransposons, which unlike their monomeric cellular counterparts (Davies 1990), are predicted to form strong homodimers. *Copia/Ty1* integrases are an exception to this overall trend, although they were shown to multimerize to form the intasome in vivo (Moore, Rinckel, and Garfinkel 1998). One possibility could be that *Copia/Ty1* intasome complexes are only formed within VLPs or in the presence of DNA substrates. In line with this hypothesis, purified *Ty1* integrases are monomeric in vitro (Nguyen et al. 2021).

Structure-based predictions also enabled us to predict multimerization between SPM-encoded TnpD and TnpA proteins, which together form a large multiprotein complex. This large complex has the potential to interact with DNA through positively charged pockets within TnpA, enabling the recruitment and homodimerization of TnpD transposases at target sequences. The long coiled-coil a-helix domains of TnpAs could also allow dimerization of complexes sitting on distinct DNA molecules, bringing them together to form the synaptosome complex. Future work assessing larger combinations of protein-DNA complexes, largely facilitated by the development of new machine learning algorithms together with Cryo-EM protein visualization, should enable the reconstruction of entire intasomes and transpososomes complexes, opening new avenues to investigate the transposition machinery.

An important outcome of our study is the discovery that TE expression is intimately intertwined within the regulatory network of cellular genes, with most TE-genes displaying idiosyncratic developmental or environmental specificities. Indeed, we found no evidence of global transcriptional reactivation of TEs in pollen, contrary to a previous report based on more limited data (Slotkin et al. 2009), and consistent with active TEs being co-expressed with developmentally or environmentally regulated genes. Furthermore, we identified numerous shared cis-regulatory sequences together with some of the TFs likely involved in their joint regulation.

While comprehensive, our functional annotation of active TEs is certainly not exhaustive, as it was based on transcriptome data obtained using specific tissues and genetic backgrounds. Similar analysis using the same or other epigenetic mutants in the reference accession as well as in strains taken from the wild, grown under different conditions will certainly expand the catalog of Arabidopsis TE-genes and products. In turn, this data may provide key information as to their collective contribution to phenotypic differences between strains and pave the way for such studies in other species.

## METHODS

### Plant materials

A. thaliana Col-0, ddm1-2, ddm1-rdr2 and ddm1-rdr6 were grown in long-days (16h:8h light:dark) at 23°C. ddm1-rdr2 and ddm1-rdr6 plants were obtained by crossing naive heterozygous ddm1 plants (i.e. plants produced after repeated backcrossing of an initial ddm1 parent) with heterozygous rdr2 or rdr6 followed by selfing and selection of F2 double mutants.

### Full-length cDNA nanopore sequencing

Total RNA was extracted from 100 mg of rosette leaves from *ddm1-2* plants using the Nucleospin RNA Plant mini kit (Macherey-Nagel). Library preparation and Nanopore sequencing were performed as previously described (Domínguez et al. 2020). Briefly, 10 ng of total RNA was amplified and converted into cDNA using SMART-Seq v4 Ultra Low Input RNA kit (Clontech). About 17 fmol of cDNA was used for library preparation using the PCR Barcoding kit (SQK-PBK004 kit, ONT) and cleaned up with 0.6× Agencourt Ampure XP beads. About 2 fmol of the purified product was amplified during 18 cycles, with a 17-min elongation step, to introduce barcodes. Samples were multiplexed in equimolar quantities to obtain 20 fmol of cDNA, and the rapid adapter ligation step was performed. Multiplexed library was loaded on an R9.4.1 flowcell (ONT) according to the manufacturer’s instructions. Standard 72-h sequencing was performed on the MinION MkIB instrument. MinKNOW software (version 19.12.5) was used for basecalling.

### Functional annotation of TE-encoding genes

Long-reads from *ddm1, ddm1rdr2, and ddm1rdr6* plants were mapped on the Arabidopsis reference genome TAIR10 using minimap v2.11-r797 (Li 2018) with the following options-ax splice-G 30k-t 12 and STAR v2.5.3a (Dobin et al. 2013) with the following options--outFilterMultimapNmax 50--outFilterMatchNmin 30--alignIntronMax 10000--alignSJoverhangMin 3, respectively. Previously published short-reads (Oberlin et al. 2017) were also mapped on the Arabidopsis reference genome (TAIR10) using STAR. Transcript annotation was performed using the FLAIR pipeline (Tang et al. 2020). First, splicing junctions based on short-read sequencing data were extracted using ‘junctions_from_sam.py’ script and used to correct ONT long-reads using ‘flair.py correct’ script. Transcript isoforms were then detected using the ‘flair.py collapse’ script and transcripts supported by at least five long-reads were retained. Exon-exon junctions were validated using publicly available short-read RNA-seq data. Raw reads were mapped on the TAIR10 genome using STAR, and ‘junctions_from_sam.py’ script was used to quantify the number of short reads supporting each junction. Transcript annotations containing at least one junction not supported by a minimum of 5 reads in total were considered to be low confidence annotations and were discarded. Transcripts sharing at least one exon-exon junction matching the TAIR10 annotation, displaying either a reciprocal coverage of over 90% with a TAIR10 annotation, or having over 75% of their length covered with them, were associated with that specific annotation. Overlapping transcript isoforms (reciprocal coverage >90%) were bundled together to the same TAIR10 entry. If more than one TAIR10 entry fulfilled the criteria, transcripts were assigned to the one with the highest overlap. Single transcripts overlapping two or more TAIR10 annotations in over 50% of the transcript length were annotated as fusions. Independent transcriptional units within a single annotation (for example, TE-genes within the same TE) were assigned as different transcripts ascribed to the same annotation but displaying a reciprocal coverage below 50%. Any remaining transcript was considered to be cryptic.

A new annotation including TAIR10 data was generated by selecting the TE-derived transcripts from our work and overlapping them with TAIR10. Other annotations (cellular genes and TAIR10-exclusive TE annotations) were directly imported from TAIR10.

### Poly-A identification

Soft-clipped bases from long-read alignments were extracted and the nucleotide content was calculated in windows of 10, 15, 20 and 25 nt from the alignment border. Reads were assigned to the forward and reverse strand when downstream or upstream ends have A or T content over 50%, respectively, in at least one of the windows. Reads with an A or T content below 50% in all windows were considered ambiguous. Annotations’s strandness were assigned as forward or reverse only if all reads produced from that annotation displayed the same orientation. Orientation was verified using publicly available stranded short-read RNAseq experiments. Briefly, the number of short-reads mapped in the forward and reverse strand was counted using samtools view, and the proportion of reads in each strand of the annotation was compared with the strand reported by our long-read pipeline and TAIR10.

### *SPM* element phylogeny

All SPM-derived proteins longer than 200 amino acids were aligned with maize TnpD and TnpA proteins (NCBI Protein Database accessions AAG17043 and AAG17044, respectively) using MAFFT 7.508 (Katoh and Standley 2013) E-INS-i algorithm. A maximum likelihood tree was built using FastTree 2.1.11 (Price, Dehal, and Arkin 2010) and each protein was identified as either TnpA or TnpD according to their phylogenetic distance to the maize proteins. Short or truncated isoforms encoding these genes were classified as TnpA-or TnpD-encoding by aligning their protein products with the products of other members of the same *SPM* family but were excluded from the phylogenetic tree.

### Characterization of *Ty1/Copia* elements with antisense transcripts

The TSS location of antisense-transcripts, defined as the read end opposite of the polyA, was searched within the TE and in the flanking 2.5 kb around the element. Antisense-transcripts for which their TSS map within the 20% terminal regions of the TE annotation were considered to be initiated at the 3’LTR. Dotplots were generated using YASS 1.16 (Noé and Kucherov 2005). Isoform expression was measured by counting the number of long reads displaying the exact same exon-exon junctions and orientations. Multiple sequence alignment of consensus reference sequences for all *Ty1/Copia* families were obtained with MAFFT E-INS-i, and a tree was built using FastTree.

### Conserved domain prediction of TE-encoded proteins

For each transcript isoform found, all putative ORFs were detected using NCBI’s standalone ORFfinder v0.4.3, restricting the start codon exclusively to ATG and requiring the encoded proteins to be at least 50 residues long. The first and the longest ORFs were selected for each transcript for subsequent analyses. Conserved domains were predicted using rpsblast 2.13 with the parameters-seg no-comp_based_stats 1-evalue 1e-5, querying the CDD Database 3.20 (Lu et al. 2020). Results were parsed using rpsbproc 0.5.

### Structural domain prediction of TE-encoded proteins

Protein structures were modeled using the local implementation of ColabFold 1.4 (Mirdita et al. 2022). The top scoring model was selected. Predicted structures for the Uniprot reference proteome of Arabidopsis UP000006548 were downloaded from https://alphafold.ebi.ac.uk/download. Structural domains were defined as groups of amino acid residues spanning over at least 50 positions and with a maximum Predicted Aligned Error of 10 Å. A global, exhaustive all-vs-all search of all the structural domains against themselves was performed using foldseek search (--cov-mode 0-a--alignment-type 1--exhaustive-search 1-e 10000), and structural domains with a local distance difference test (LDDT) score of 90% and an alignment covering 90% of both domains were clustered together.

All SDs of each cluster were aligned against the RCSB, AlphaFold2-modeled Swissprot databases (downloaded using foldseek utilities), and the Uniprot reference proteome of Arabidopsis UP000006548 using foldseek search (--cov-mode 2-c 0.8-a--alignment-type 1-e 0.01), selecting hits covering at least 80% of the SD length and an e-value below 0.01. A main name was manually assigned for each cluster based on the best-scoring hits with previously reported structures (RCSB, Swissprot or Uniprot) and CDs found among its members.

Then, for each cluster with no assigned name, the medoid structure was calculated and aligned against all clusters with functions assigned using ChimeraX’s matchmaker implementation. If an alignment with an RMSD below 3 Å over two thirds of the SD length was found, the unknown cluster was renamed as the matching named cluster.

### Chromosome slippage RNA secondary structure

The putative slippage site of *AT1TE42395* was located between the last stop codons of the 0 frame and the next following stop codon at the-1 frame. The RNA secondary structure was predicted using mfold (Zuker 2003), limiting loop formation to bases at most 25bp away and using 0 as the window parameter.

### Integrase C-terminal domain characterization

*Ty1/Copia* elements containing integrase and retrotranscriptase SDs, and Ty3/Gypsy elements containing full-length integrase, were selected. For Ty1/Copia elements, the linker region connecting both SDs was scanned for protease cleavage sites based on residue hydrophobicity index Hi (Kyte and Doolittle 1982), using sliding windows of 4 residues (p2, p1, p1’ and p2’). The sum of differences in hydrophobicity between these sites and reported cleavage site sequences (Merkulov et al. 1996; Kirchner and Sandmeyer 1993) was used to score each window, and the best scoring window on each linker was selected as the most likely protease cutting site. For Ty3/Gypsy elements, the last 200 residues of the polyprotein, up to the end of the integrase SD, were selected. These sequences were then scanned for putative nuclear localization signals. A gapped motif for these sequences was predicted using GLAM2 with the options-r 20-n 100000-a 2-b 20.

### Structure-based dimerization and DNA-binding potential prediction

Protein dimers were modeled using AlphaFold-Multimer (Evans et al. 2022) and dimerization potential was determined based on interatomic distance. Residues from one chain at an interatomic distance below 8 Å between the β-carbons (α-carbon in the case of glycine residues) from any residue on the other chain were defined as interacting residues. Pairs of proteins with at least 10 interacting residues on each chain were considered to form dimers. To model the complete transpososome, the different possible trimers were modeled and the complete transpososome was assembled by aligning these trimers using ChimeraX matchmaker. To identify putative DNA-binding pockets, positively charged protein’s surface were identified using the APBS algorithm implemented in ChimeraX. DNA-binding residues were confirmed using GraphSite (Yuan et al. 2022).

### Analysis of publicly available transcriptome dataset

462 transcriptome datasets were retrieved from the Sequence Read Archive and the European Nucleotide Archive (Supplementary Table 1). For each sample, reads were mapped using STAR, discarding reads with less than 30 matching bases and with a maximum intron length allowed of 10kb. PCR duplicates were removed using Picard 1.99 MarkDuplicates. Read counts were generated for all the features in the combined TAIR10 + TE-gene annotation using featureCounts (Liao, Smyth, and Shi 2014), assigning fractional counts to multimapping reads and to those spanning over more than one feature (-O-M--fraction). Features with a coverage below 10 reads in all samples were filtered out, and samples with a total coverage of less than 3 million mapped reads were discarded. Read counts were normalized by size factors and then transformed using the variance stabilizing transformation implemented in the R package DESeq2 (Love, Huber, and Anders 2014) using estimateSizeFactors and vst. Variance stabilized counts used for subsequent analyses. Batch effects for different sequencing experiments were controlled using the R package ComBat.

### Coexpression network analysis

A signed weighted gene coexpression network was built using the R package wgcna (Langfelder and Horvath 2008) with a β coefficient of 10. Coexpression clusters were defined using the function blockwiseModules. GO term enrichment for each cluster was performed using the R package clusterProfiler (Yu et al. 2012). For visualization, the complete topological overlap matrix was pruned to keep only the 10 edges with the highest weight for each node and plotted using Cytoscape (Shannon et al. 2003). Expression of TE-genes located closer than 1kb away from a coexpressed cellular gene was considered to be driven by positional effects, and these genes were hidden from the network. Full-length copies of *ATCOPIA78* elements were defined as copies longer than 90% of the length of the autonomous *AT5TE15240*. Copies below this size were designated as truncated.

### Analysis of DAP-seq data

DNA affinity purification and sequencing (DAP-seq) data obtained in Arabidopsis thaliana (O’Malley et al. 2016) for 529 TFs were processed using a modified version of the bioinformatics pipeline implemented before (O’Malley et al. 2016). Specifically, reads mapping to repetitive sequences were also considered to detect binding over TEs. Reads were mapped on the TAIR10 genome using Bowtie2 v.2.3.2 (Langmead and Salzberg 2012), and PCR duplicates were removed using Picard. The detection of peaks for TF binding was performed with GEM (Guo, Mahony, and Gifford 2012), arguments--k_min 6--kmax 20--k_seqs 600--k_neg_dinu_shuffle--t 5. TFBS within DAP-seq peaks were detected with FIMO (Grant, Bailey, and Noble 2011) using the corresponding TF motifs (O’Malley et al. 2016). Hits with a reciprocal coverage over 80% were merged, with a maximum length dictated by the motif length as retrieved from the database. Only DAP-seq peaks containing cognate TF binding motifs were considered. Relative distance between TFBS and the closest TSS was measured for each TE-gene and cellular gene from the center of the DAP-seq peak to the exact TSS position extracted from the transcript coordinates and orientation. To calculate the distance distribution of TFBS from coexpressed and non-coexpressed TFs, only the TFBS found within the TE boundaries were considered, and only TE-TF pairs on the same coexpression modules were marked as coexpressed. Significance was assessed by performing independent Mann-Whitney U tests per TE family and using the Benjamini-Hochberg correction for multiple testing.

## Authors’s contributions

LQ conceived the project. CB performed most bioinformatic analysis, interpreted the data, and wrote the manuscript. BL performed initial transcriptional network analysis. VC interpreted the data and revised the manuscript. LQ performed ONT transcriptome experiments, interpreted the data, and wrote the manuscript. All the authors read and approved the paper.

## Acknowledgements

We thank members of the Quadrana and Colot groups for discussions. We thank especially Erwann Caillieux for providing the *ddm1rdr2* and *ddm1rdr6* mutant lines as well as for assistance. We also thank Taku Sasaki for critical reading of the manuscript. This work was supported by the European Research Council (ERC) under the European Union’s Horizon 2020 research and innovation program (grant agreement No. 948674 to LQ).

## Availability of data and materials

The datasets generated during the current study and cDNA sequences of full-length TE transcripts are available in the European Nucleotide Archive (ENA) repository PRJEBXXX. Annotations of TE gene and encoded transcripts, putative TE-encoded protein sequences and structures, and gene regulatory networks are publicly available at Zenodo (doi:10.5281/zenodo.10831665), as well as available as supplementary material.

---

**Figure S1.**
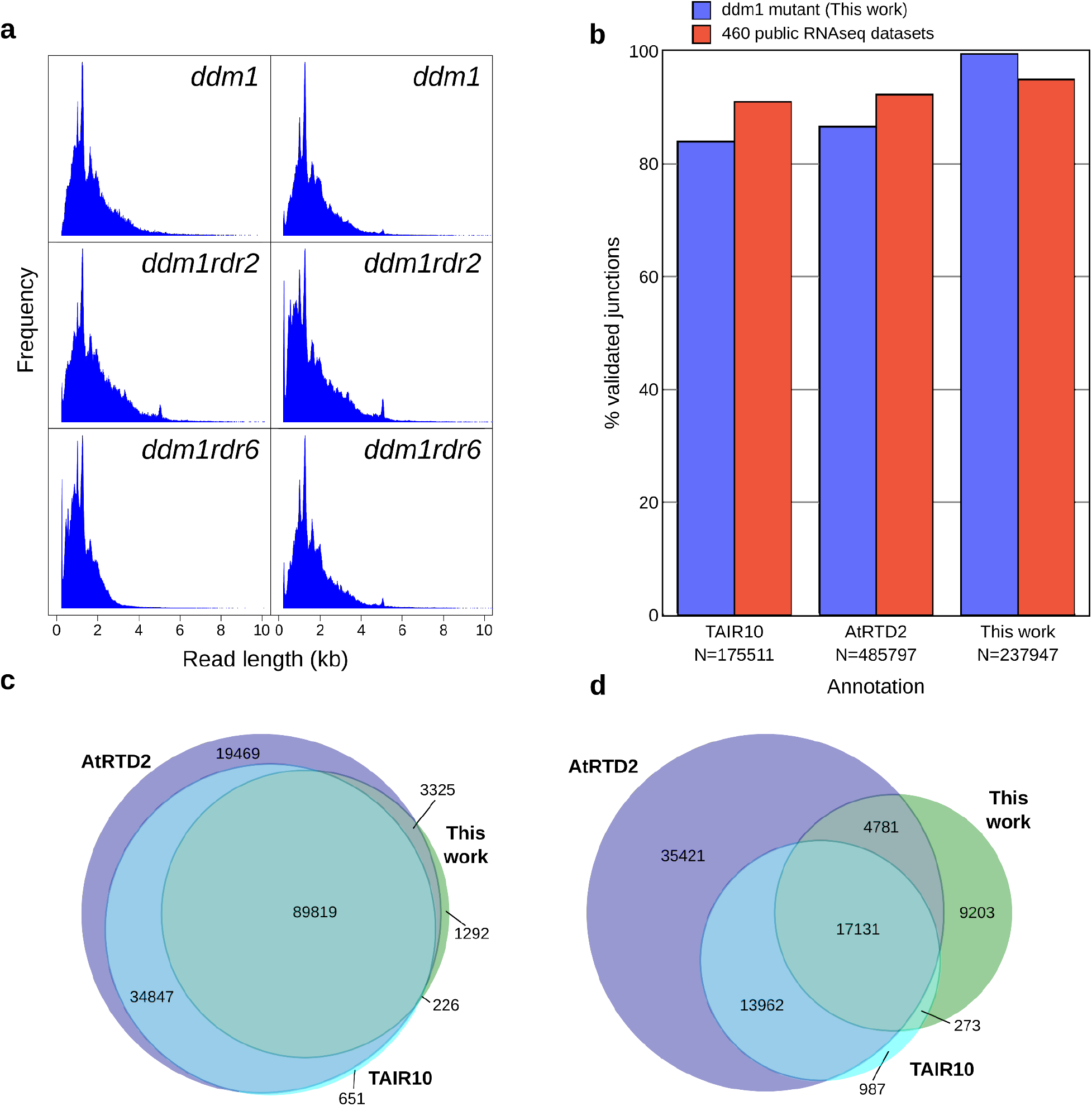
Validation of annotated junctions. **a**. Read length distribution of the different ONT sequencing experiments. **b**. Percentage of exon-exon junctions supported by at least five short reads in either the *ddm1* transcriptome (Oberlin et al. 2017) or in the public transcriptomic data detailed in Supplementary table 1. The total number of annotated junctions N is included for each annotation. **c**. Venn diagram of private and shared exon-exon junctions among AtRTD2, TAIR10 and this work. **d**. Venn diagram of private and shared isoforms among AtRTD2, TAIR10 and this work. Each isoform is defined as a unique set of exon-exon junctions.

**Figure S2:**
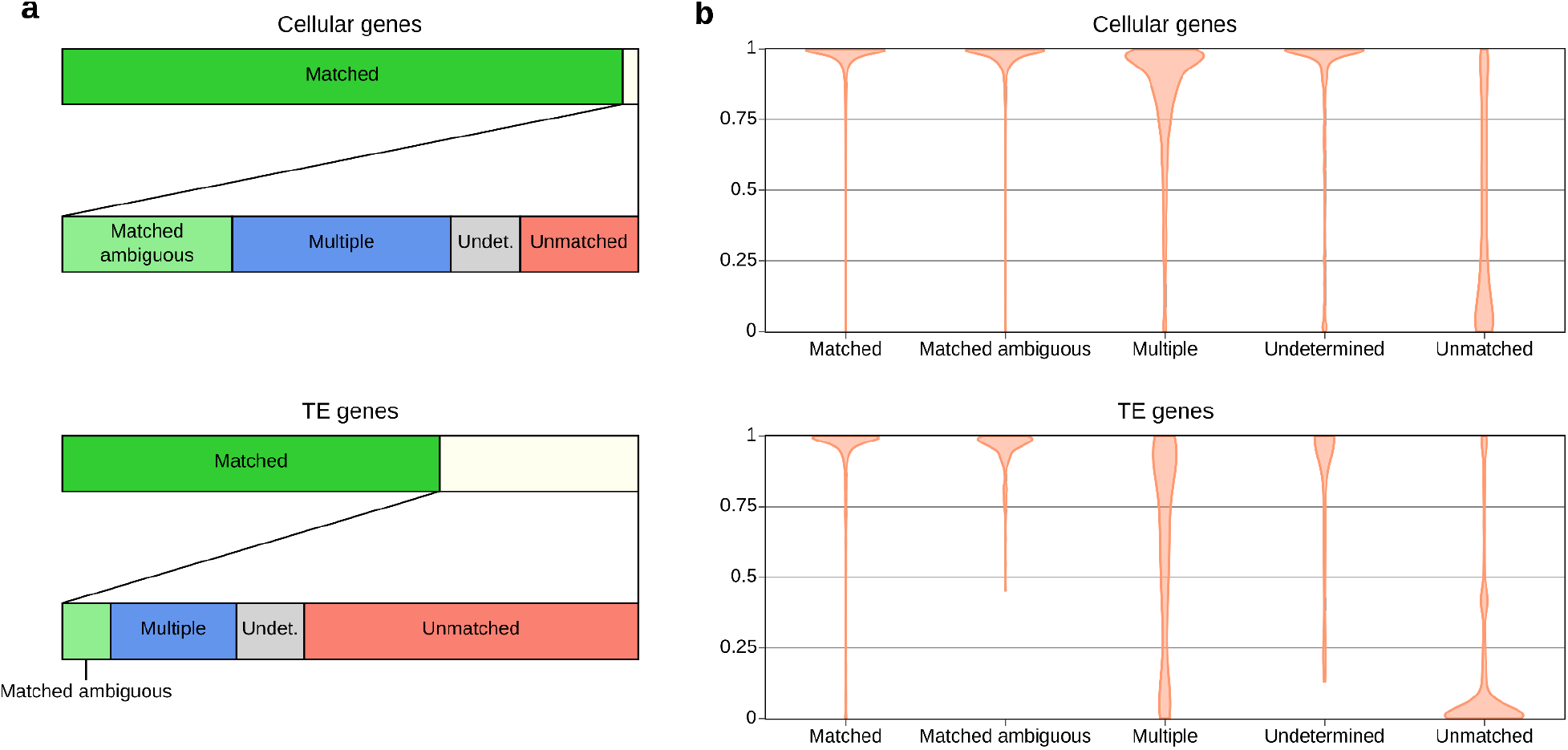
Strandness comparison between TAIR10 and this work. **a.** Strand assignment comparison between TAIR10 and our poly-A detection pipeline for genes (top) and TE-genes (bottom) covered by long-read sequencing. ONT-annotated features with isoforms with both matching and undetermined orientations are labeled as “matched ambiguous”. ONT-annotated features with isoforms displaying conflicting strands are labeled as “multiple”. ONT-annotated features with no evidence for any orientation are labeled as “undetermined”. Only ONT-annotated features with isoforms displaying an orientation reverse to TAIR10 are labeled as “unmatched”. **b**. Distribution of the number of short reads from 46 stranded RNA-seq public datasets (Supplementary Table 1) supporting the orientation reported by TAIR10 in cellular and TE-genes.

**Figure S3:**
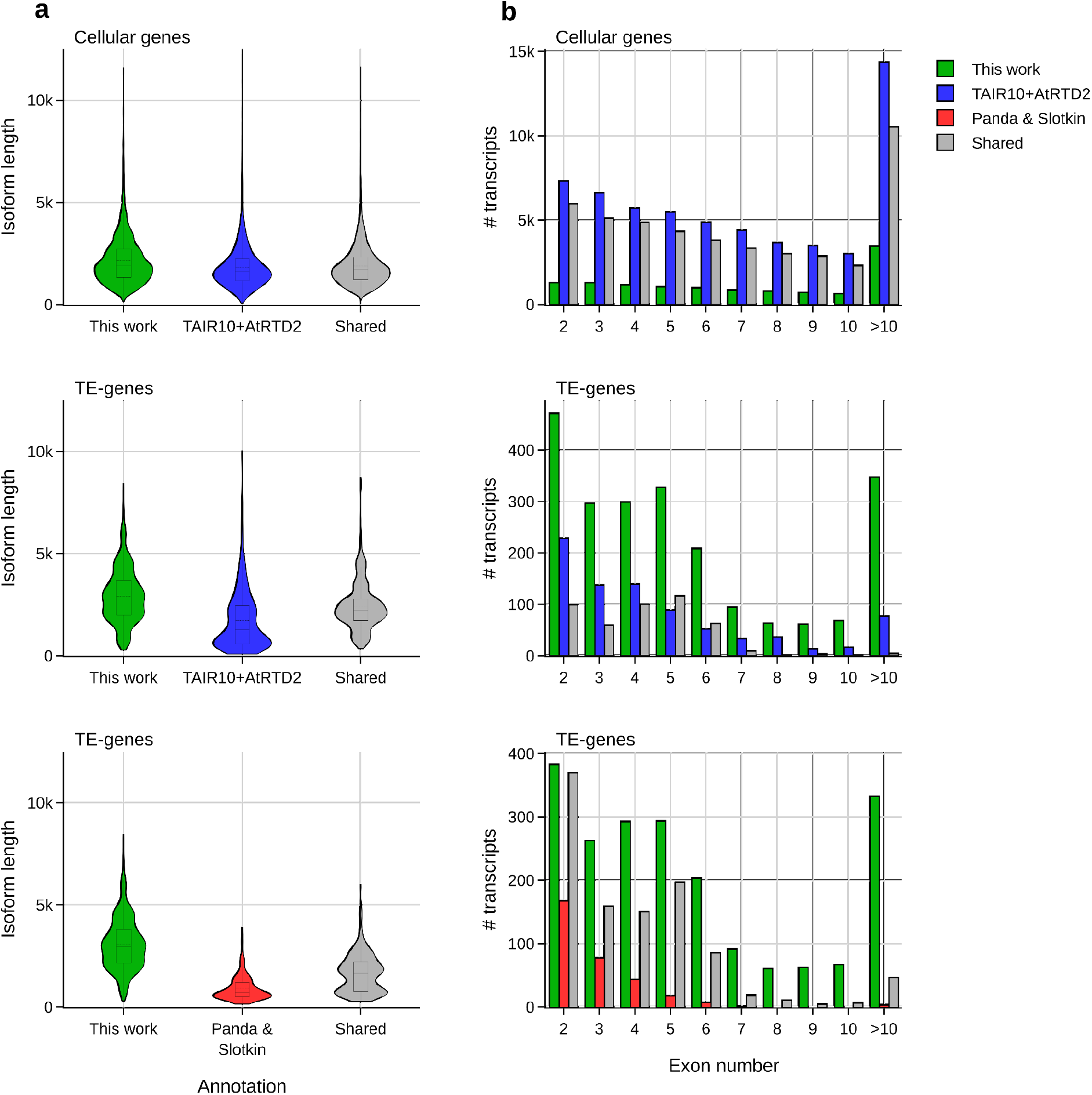
Annotation comparison. **a**. Length distribution of each private or shared isoform between this work and prior annotations for genes (top) and TEs of TAIR10+AtRTD2 (mid) or the TEs from the prior study of Panda and Slotkin (bottom). **b**. Exon count distribution of multiple exon transcripts for exclusive and shared isoforms described by this work, TAIR10+AtRTD2 or Panda and Slotkin work.

**Figure S4.**
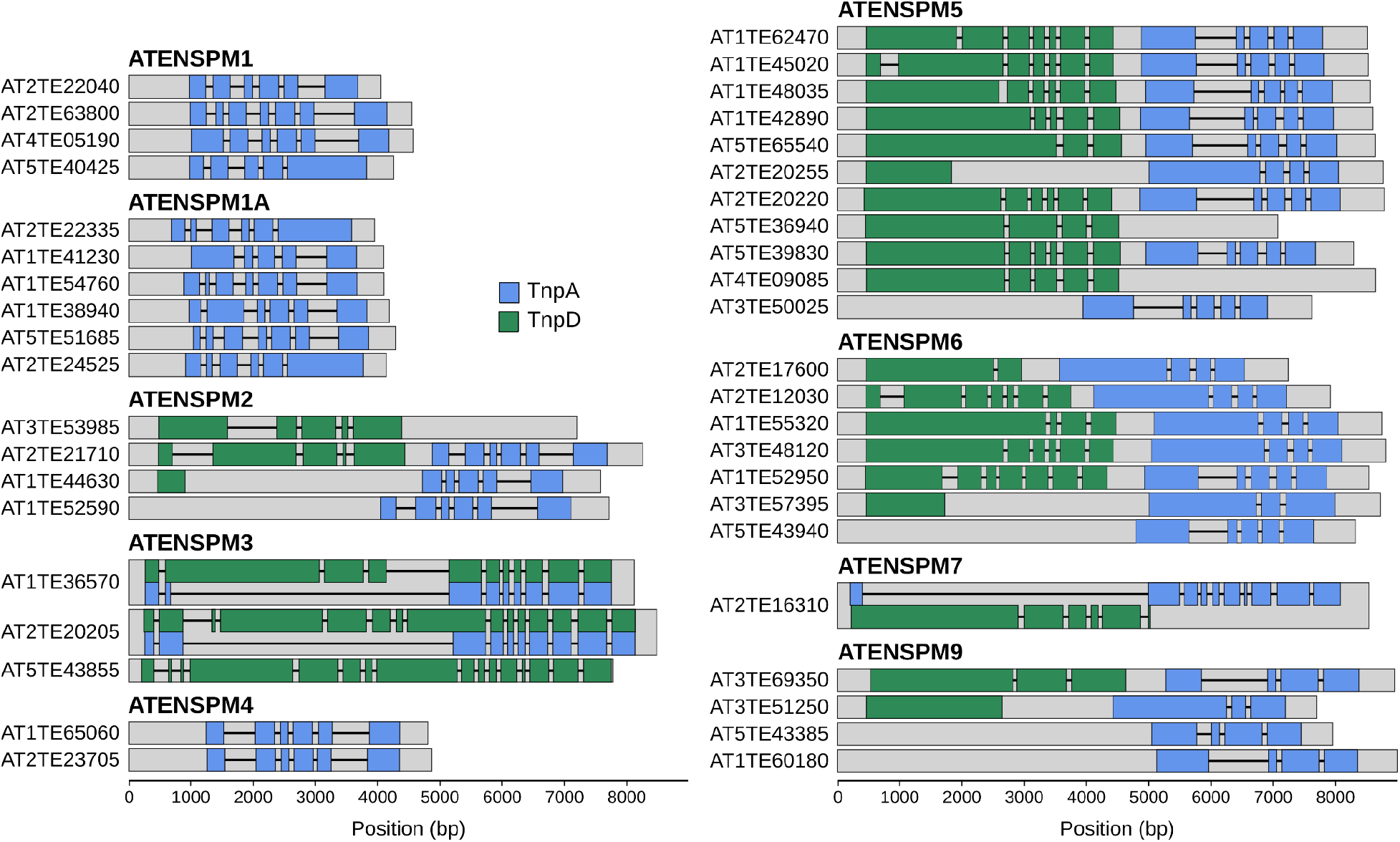
*SPM* architectures on the *Arabidopsis thaliana* genome. TE-genes and isoforms from all the *SPM* elements characterized in this work. The identity of each transcript was assigned by aligning the proteins encoded by them with the *Zea mays* TnpA and TnpD (NCBI protein database accessions AAG17044 and AAG17043, respectively).

**Figure S5.**
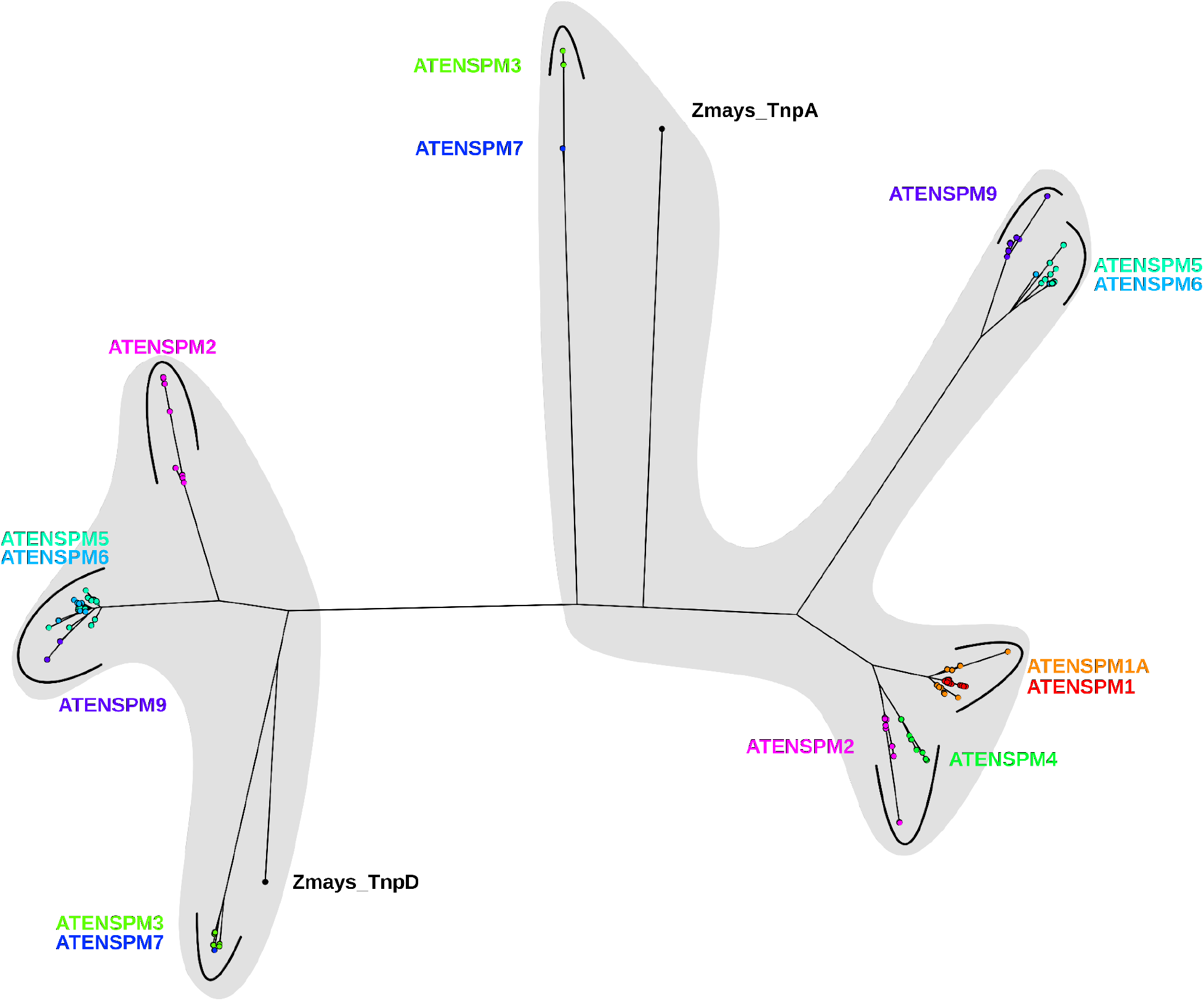
Phylogenetic tree of *SPM*-encoded proteins. All proteins with a length over 200 amino acids encoded by transcripts derived from *SPM1 to SPM9* elements are included. Two major clades are highlighted.

**Figure S6.**
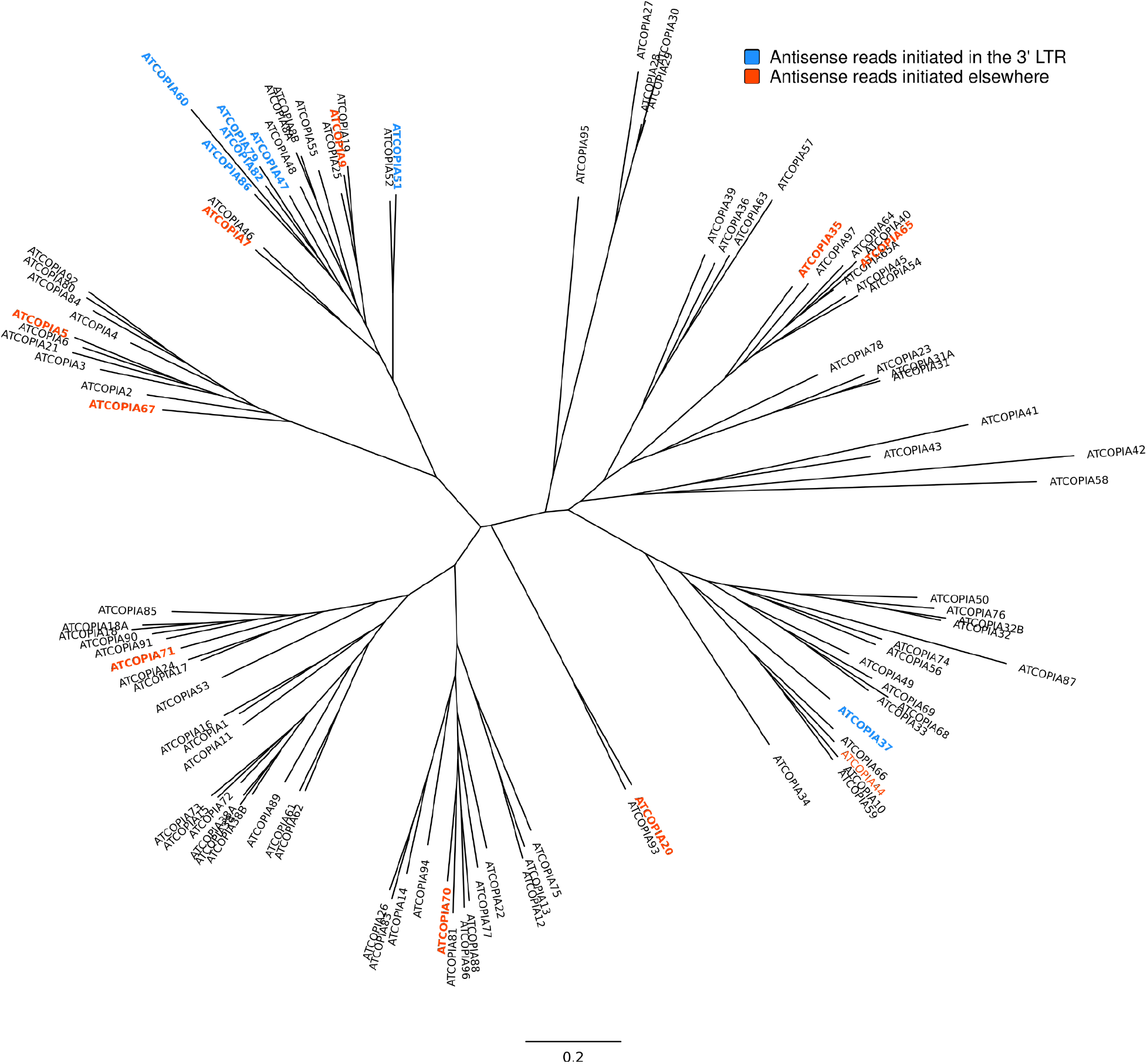
Phylogenetic tree of *Ty1/Copia* elements. The consensus sequences of each *Ty1/Copia* family were used to build the tree. Families with antisense transcription not generated in the 3’LTR are shown in red. Families with elements harboring 3’LTR-generated antisense, polyexonic transcripts are shown in blue. For the remaining families, no antisense transcription was detected.

**Figure S7.**
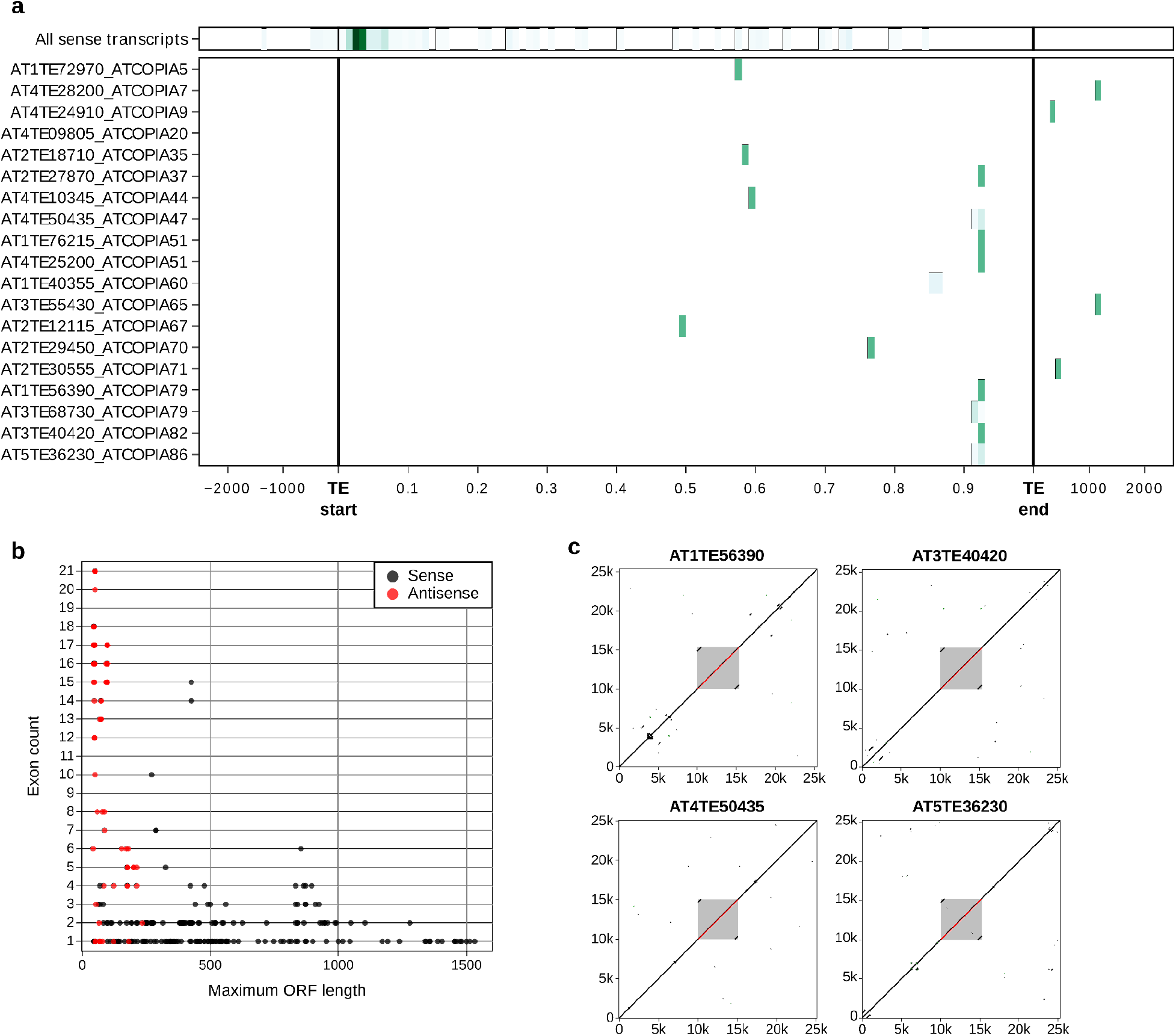
Characterization of Copia antisense transcripts. **a**. Transcription start site for all sense (top) and antisense (bottom) Copia transcripts. **b**. Protein length (in residues) of the longest ORF per transcript for all Copia elements. Antisense transcripts are marked in red. **c**. Dotplot of selected Copia elements generating antisense transcripts and 10kb of their flanking regions. Each element is highlighted in red, and its span is shown as a grey box.

**Figure S8.**
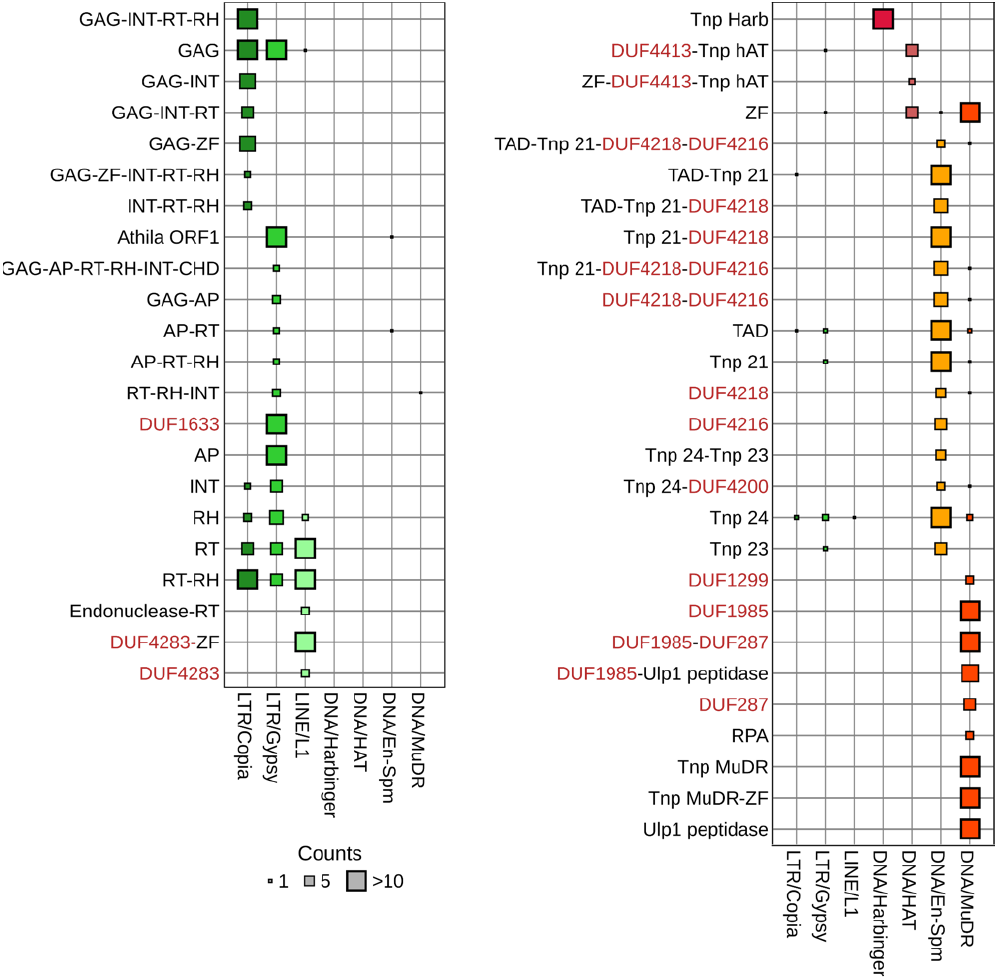
Conserved domain architectures. Main conserved domain architectures for Class I (left) and Class II (right). Point size indicates the number of TE elements harboring at least one protein with a given architecture. Domains of Unknown functions (DUF) are highlighted in red.

**Figure S9.**
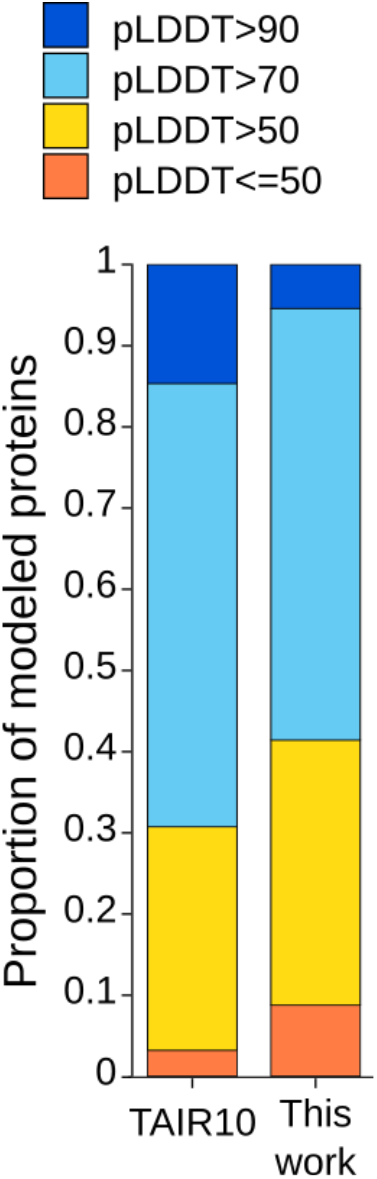
pLDDT distribution. pLDDT distribution of modeled TE-encoded proteins and the Arabidopsis reference proteome available at AlphaFold Protein Structure Database.

**Figure S10.**
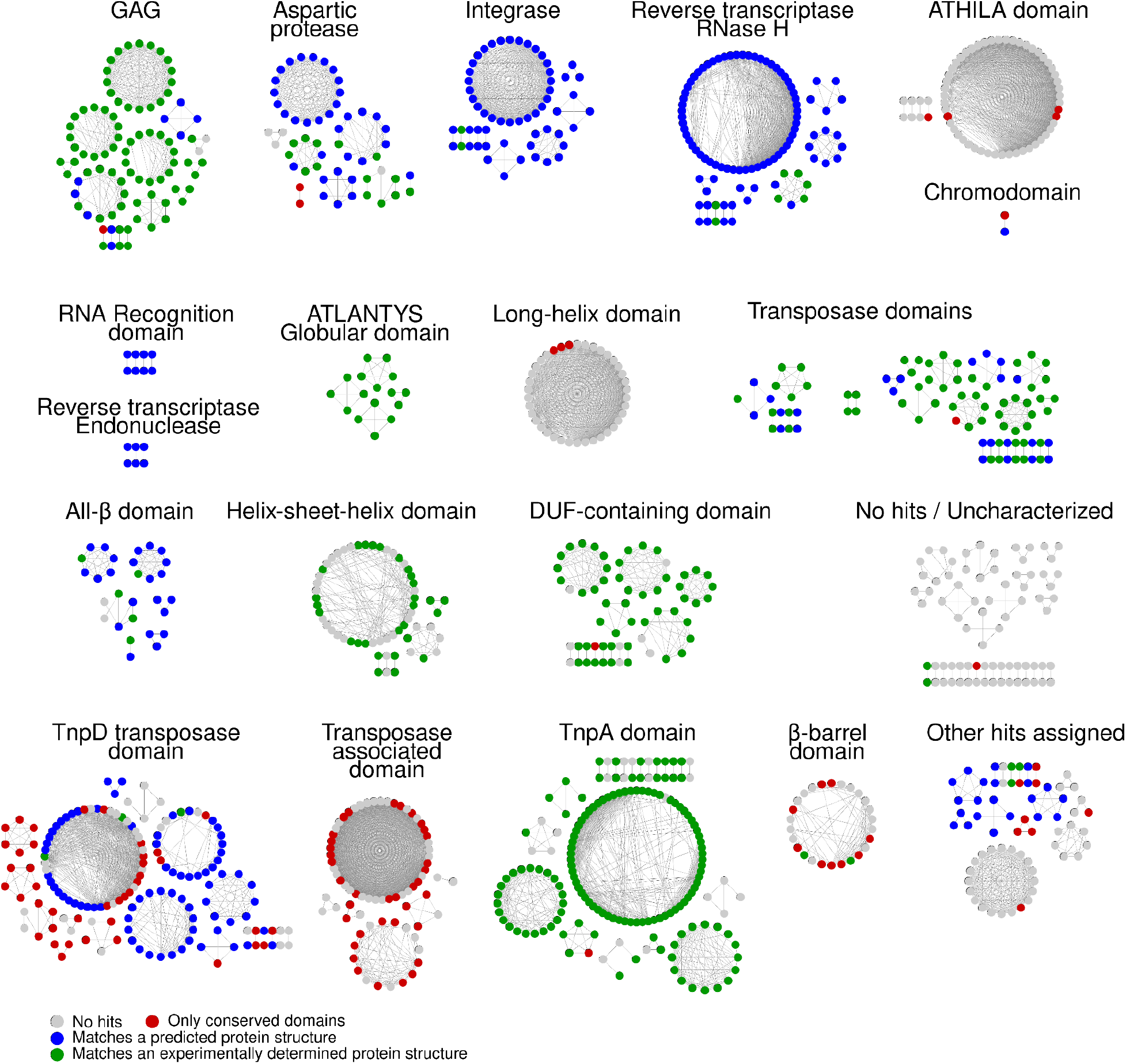
Structural domain network. Nodes represent SDs, and connecting lines indicate structural similarity between SDs. Nodes are colored based on their sequence or structural match with previously characterized proteins. Only clusters grouping at least two SDs are shown.

**Figure S11.**
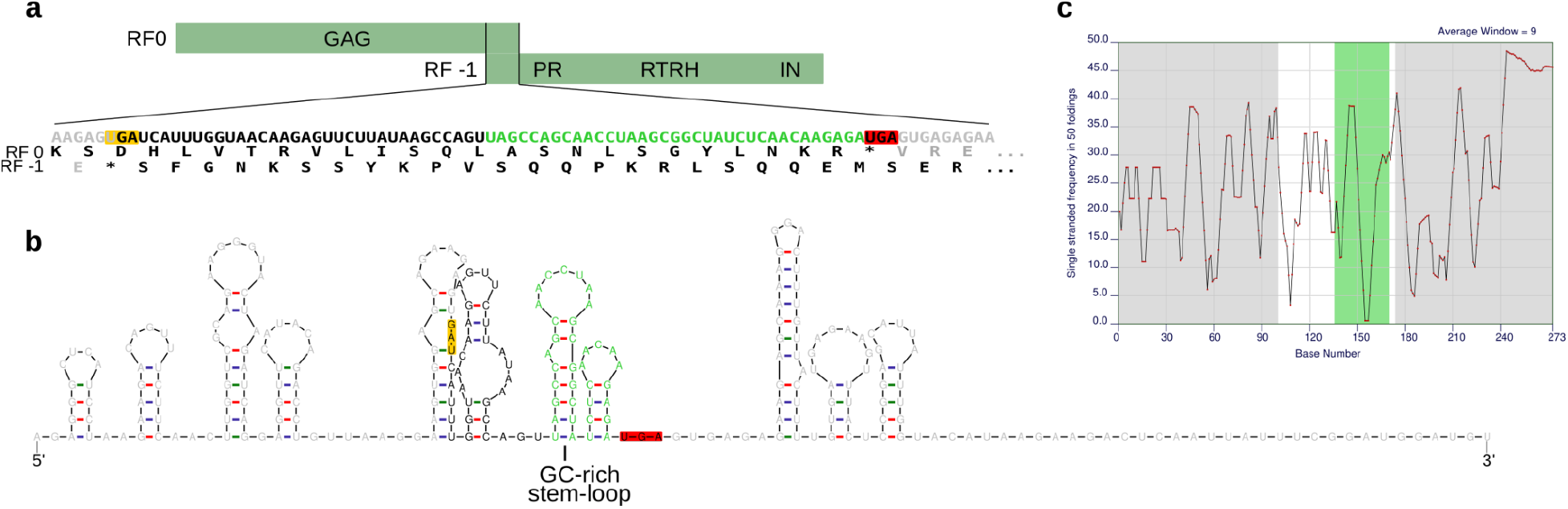
Ribosomal slippage site of *ATGP3*. **a**. Schematic representation of the frameshift location on *ATGP3* (*AT1TE42395*). The sequence where the slippage should occur for a single uninterrupted ORF to be maintained is shown below. The first downstream and upstream stop codons on the 0 and-1 reading frames are highlighted in red and orange, respectively. **b**. Minimal energy predicted RNA secondary structure along the slippage region extended by 100bp. Extended bases are shown in grey and the stop codons are highlighted in red and yellow. 36 bases forming two stem-loops preceding the downstream stop codon are shown in green. **c**. Frequency of single strand RNA secondary structure along the slippage region across 50 minimal energy structures. The green and grey areas mark the bases outside the slippage region and the two loops.

**Figure S12.**
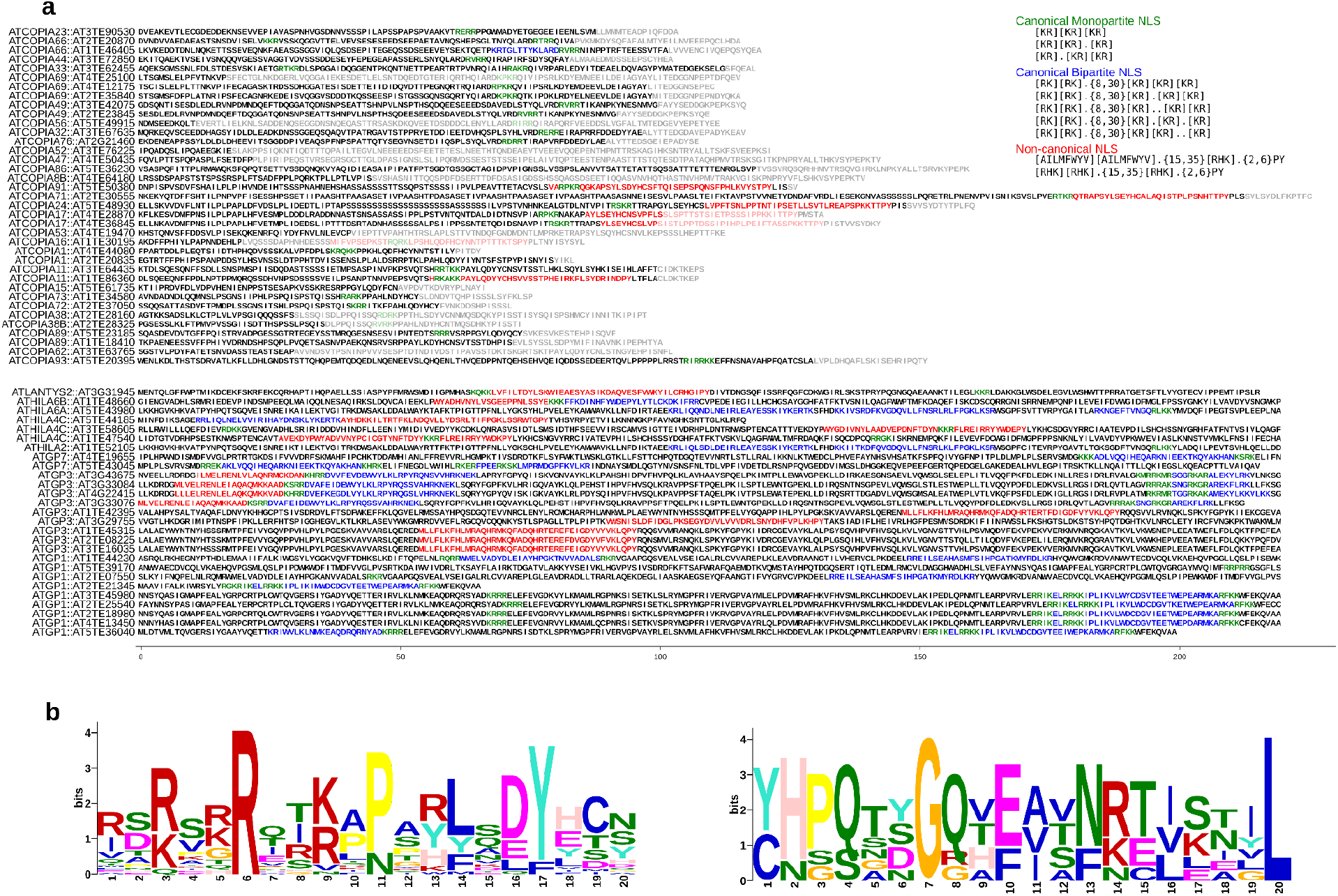
Nuclear localization signals in integrase C-terminal domains. a. Disorganized C-terminal domain of Ty1/Copia (top) and Ty3/Gypsy (bottom) elements. For Gypsy elements, the chromodomain is also included. Monopartite, bipartite and non-canonical NLS are shown in green, blue and red. **b**. Motif detected for Ty1/Copia (left) and Ty3/Gypsy (right) elements. A single element per family was used for motif construction.

**Figure S13.**
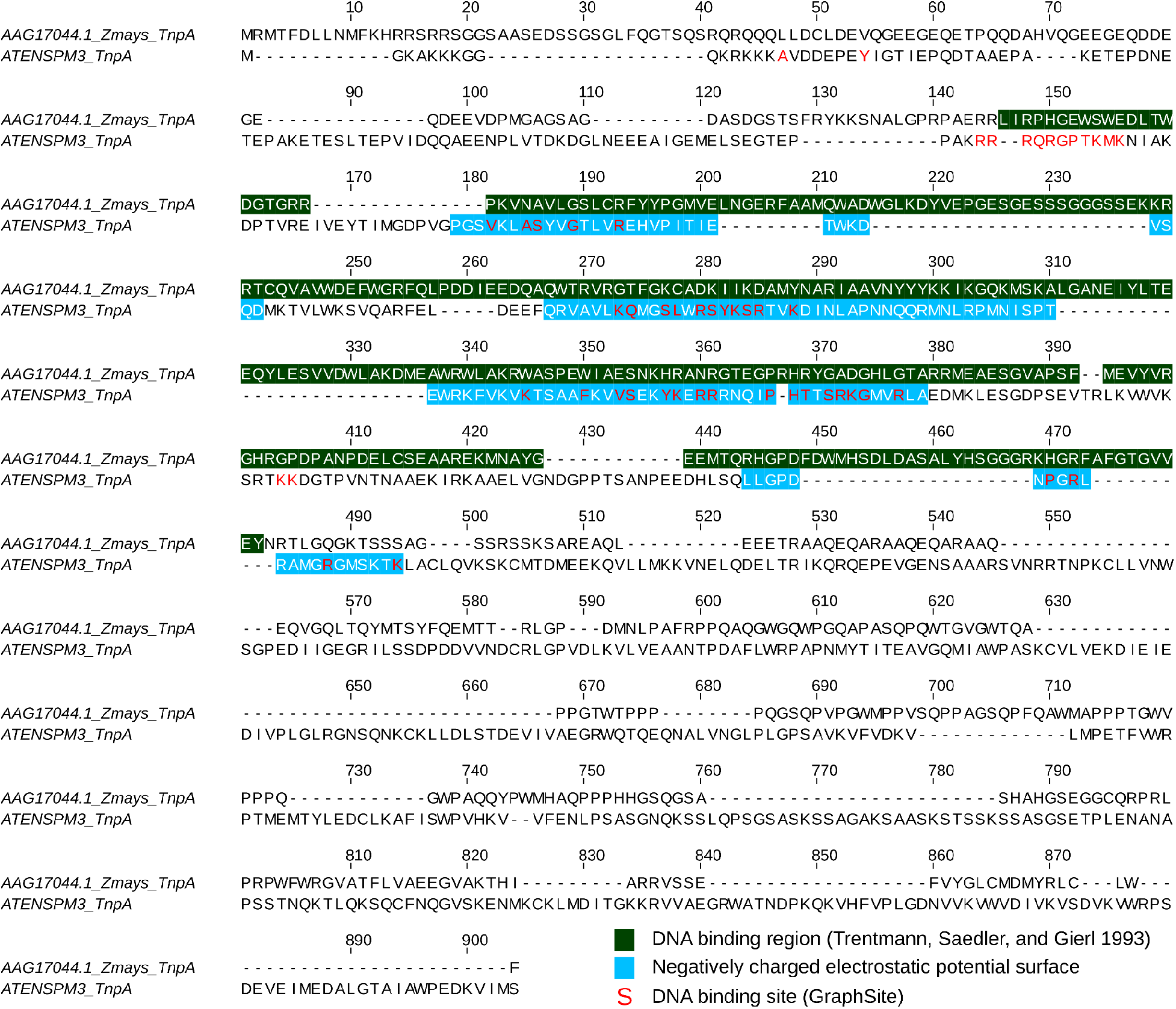
Identified DNA-binding sites from *SPM* TnpA. Protein alignment of the modeled TnpA from *ATENSPM3* (AT2TE20205) and a previously reported *Zea mays* TnpA (AAG17044). DNA binding regions predicted by alternative methods are shown.

**Figure S14.**
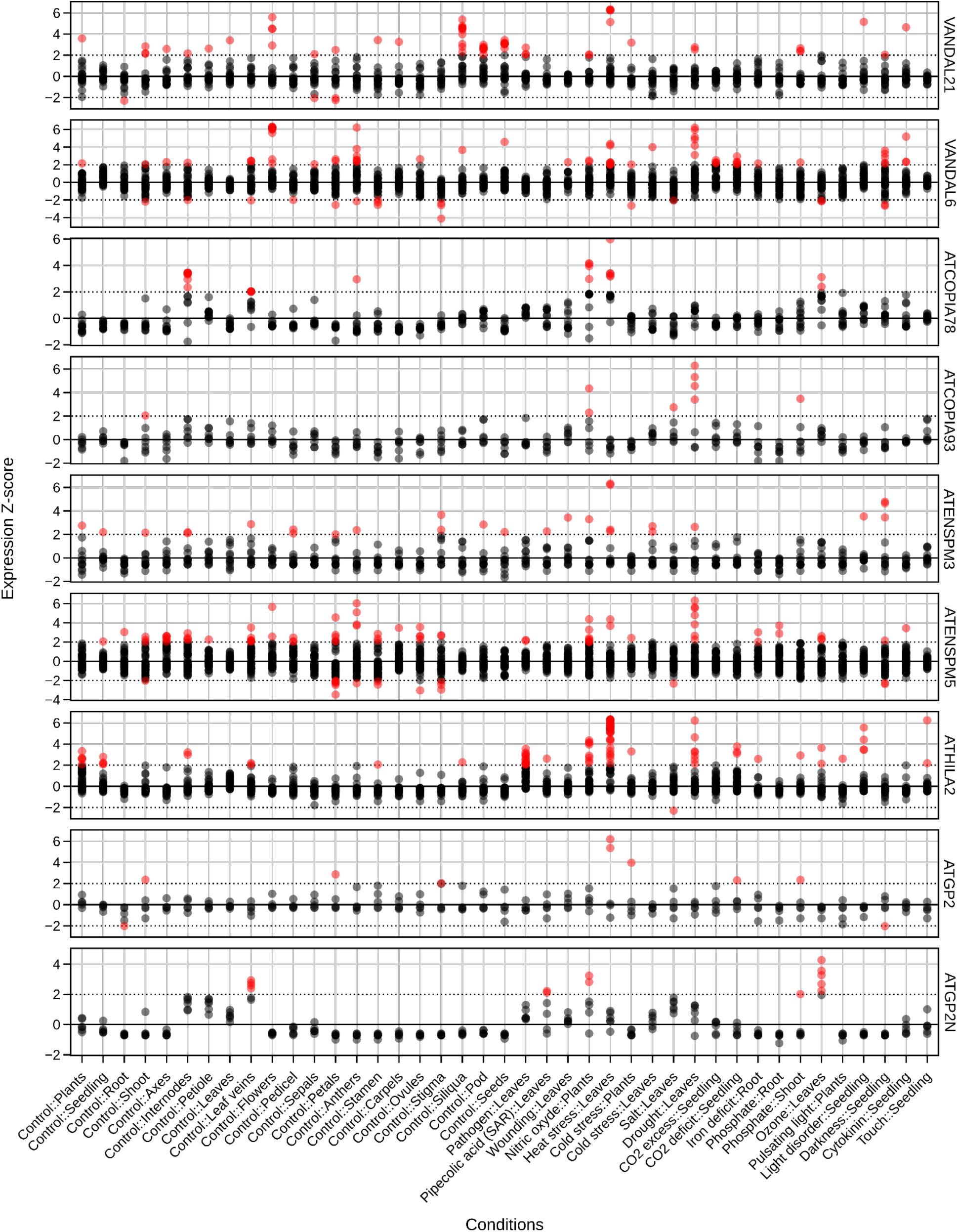
Expression of selected TEs over an array of tissues and conditions. Average normalized expression of all TE-genes from selected TEs across transcriptomic experiments. Z-scores were calculated for each TE-gene and condition compared with the expression of the same gene in all conditions. Z-score values above 2 are highlighted in red.

**Figure S15.**
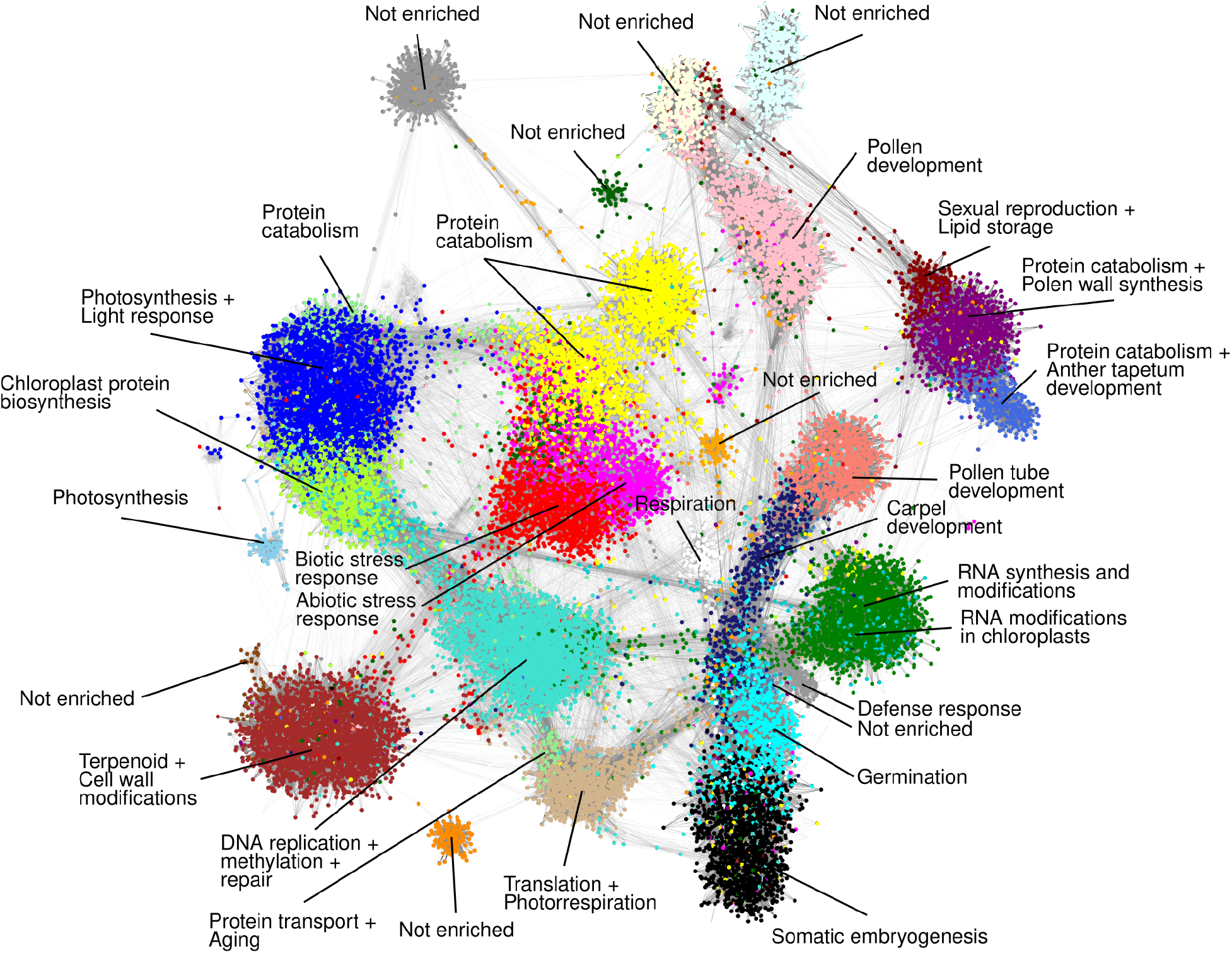
Coexpression clusters and their associated functions. Coexpression clusters are shown in different colors. Genes not included in any cluster are hidden. Major GO terms associated with each cluster are shown, for the full list see Supplementary Table 3.

**Figure S16.**
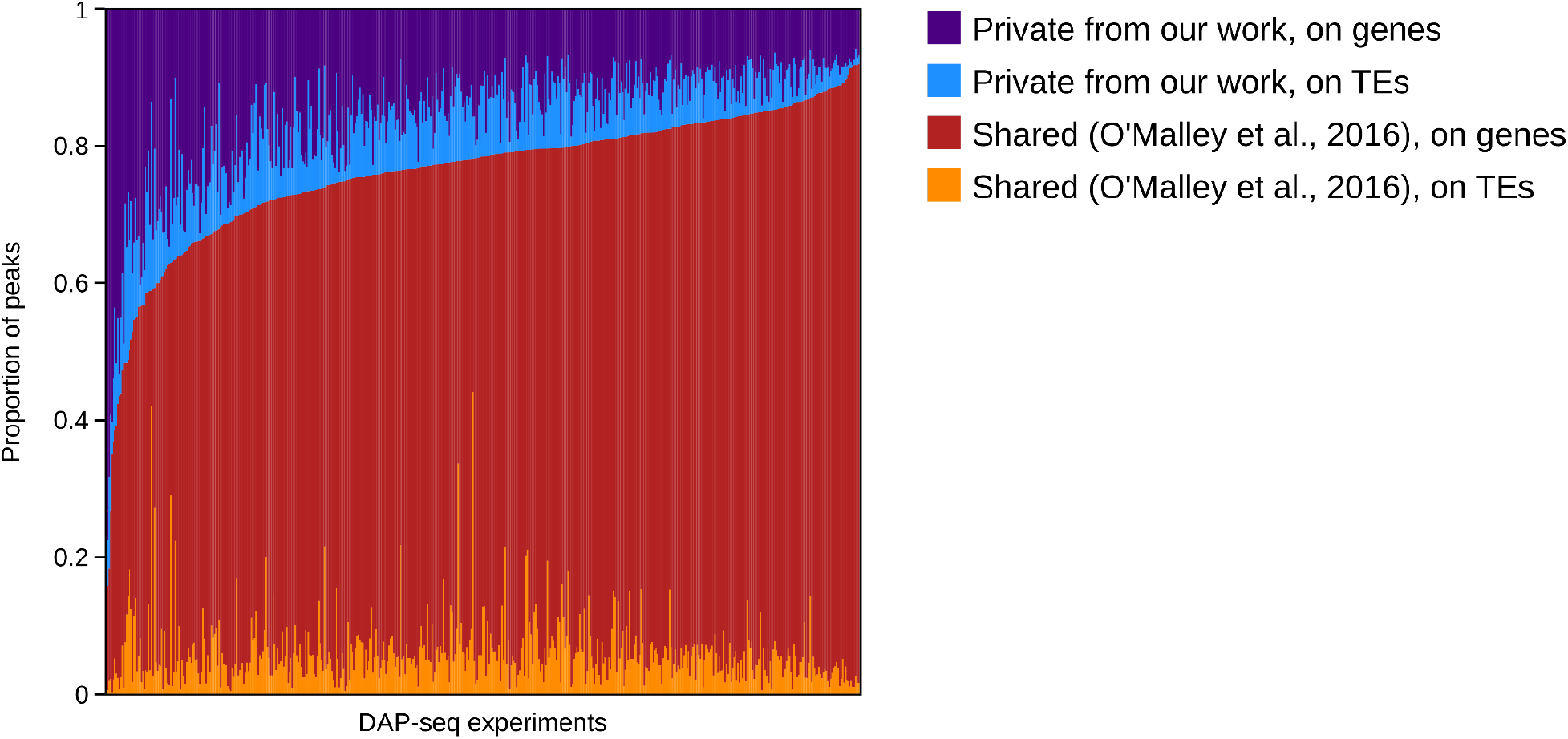
DAP-seq peak calling. Proportion of DAP-seq peaks shared between our analysis and that reported previously (O’Malley et al 2016).

**Figure S17.**
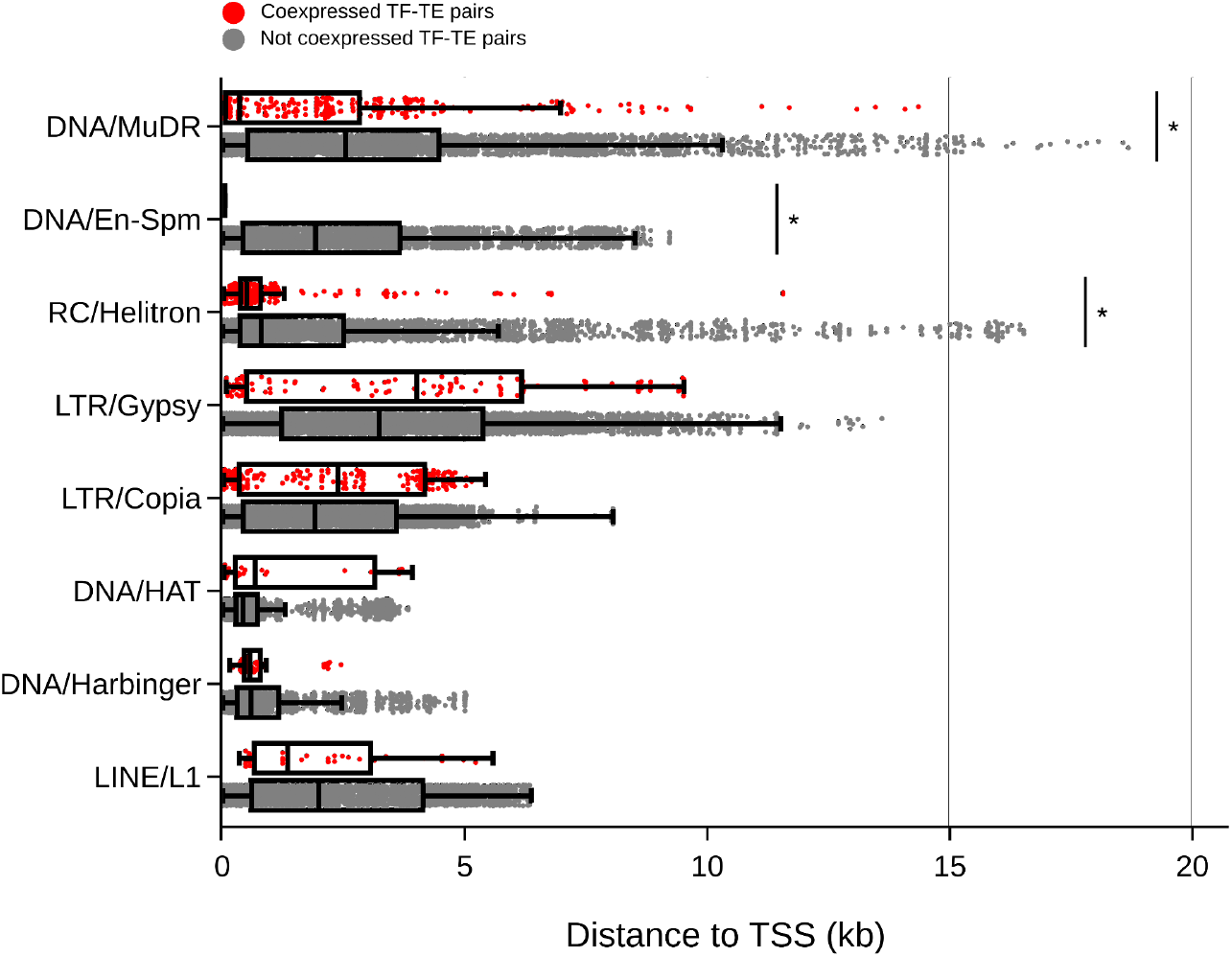
Distribution of TFBS distances to TSS for coexpressed and non-coexpressed TF– TE pairs. Distance between each TFBS overlapping a TE and the TSS of all TE-genes within the element. Coexpressed pairs of TFs and TE-genes are shown in red, while non-coexpressed pairs are shown in gray. Asterisks mark statistically significant different distributions between coexpressed and non-coexpressed pairs (Mann-Whitney U test + Benjamini-Hochberg correction, p-value < 0.05).

## REFERENCES

1. Abascal-Palacios, Guillermo, Laura Jochem, Carlos Pla-Prats, Fabienne Beuron, and Alessandro Vannini. 2021. “Structural Basis of Ty3 Retrotransposon Integration at RNA Polymerase III-Transcribed Genes.” Nature Communications 12 (1): 6992.

2. Armstrong, Susan J., Anthony P. Caryl, Gareth H. Jones, and F. Christopher H. Franklin. 2002. “Asy1, a Protein Required for Meiotic Chromosome Synapsis, Localizes to Axis-Associated Chromatin in Arabidopsis and Brassica.” Journal of Cell Science 115 (Pt 18): 3645–55.

3. Asif-Laidin, Amna, Christine Conesa, Amandine Bonnet, Camille Grison, Indranil Adhya, Rachid Menouni, Hélène Fayol, Noé Palmic, Joël Acker, and Pascale Lesage. 2020. “A Small Targeting Domain in Ty1 Integrase Is Sufficient to Direct Retrotransposon Integration Upstream of tRNA Genes.” The EMBO Journal 39 (17): e104337.

4. Aziz, Ramy K., Mya Breitbart, and Robert A. Edwards. 2010. “Transposases Are the Most Abundant, Most Ubiquitous Genes in Nature.” Nucleic Acids Research 38 (13): 4207–17.

5. Baduel, Pierre, Basile Leduque, Amandine Ignace, Isabelle Gy, José Gil Jr, Olivier Loudet, Vincent Colot, and Leandro Quadrana. 2021. “Genetic and Environmental Modulation of Transposition Shapes the Evolutionary Potential of Arabidopsis Thaliana.” Genome Biology 22 (1): 138.

6. Barro-Trastoy, Daniela, and Claudia Köhler. 2024. “Helitrons: Genomic Parasites That Generate Developmental Novelties.” *Trends in Genetics: TIG*, February. 10.1016/j.tig.2024.02.002.

7. Belcourt, M. F., and P. J. Farabaugh. 1990. “Ribosomal Frameshifting in the Yeast Retrotransposon Ty: tRNAs Induce Slippage on a 7 Nucleotide Minimal Site.” Cell 62 (2): 339–52.

8. Berthelier, Jérémy, Leonardo Furci, Shuta Asai, Munissa Sadykova, Tomoe Shimazaki, Ken Shirasu, and Hidetoshi Saze. 2023. “Long-Read Direct RNA Sequencing Reveals Epigenetic Regulation of Chimeric Gene-Transposon Transcripts in Arabidopsis Thaliana.” Nature Communications 14 (1): 3248.

9. Chuong, Edward B., Nels C. Elde, and Cédric Feschotte. 2016. “Regulatory Evolution of Innate Immunity through Co-Option of Endogenous Retroviruses.” Science 351 (6277): 1083–87.

10. Davies, D. R. 1990. “The Structure and Function of the Aspartic Proteinases.” Annual Review of Biophysics and Biophysical Chemistry 19: 189–215.

11. Dobin, Alexander, Carrie A. Davis, Felix Schlesinger, Jorg Drenkow, Chris Zaleski, Sonali Jha, Philippe Batut, Mark Chaisson, and Thomas R. Gingeras. 2013. “STAR: Ultrafast Universal RNA-Seq Aligner.” Bioinformatics 29 (1): 15–21.

12. Domínguez, Marisol, Elise Dugas, Médine Benchouaia, Basile Leduque, José M. Jiménez-Gómez, Vincent Colot, and Leandro Quadrana. 2020. “The Impact of Transposable Elements on Tomato Diversity.” Nature Communications 11 (1): 4058.

13. Evans, Richard, Michael O’Neill, Alexander Pritzel, Natasha Antropova, Andrew Senior, Tim Green, Augustin Žídek, et al. 2022. “Protein Complex Prediction with AlphaFold-Multimer.” bioRxiv. 10.1101/2021.10.04.463034.

14. Fedoroff, N. V. 1989. “About Maize Transposable Elements and Development.” Cell 56 (2): 181–91.

15. Fu, Yu, Akira Kawabe, Mathilde Etcheverry, Tasuku Ito, Atsushi Toyoda, Asao Fujiyama, Vincent Colot, Yoshiaki Tarutani, and Tetsuji Kakutani. 2013. “Mobilization of a Plant Transposon by Expression of the Transposon-Encoded Anti-Silencing Factor.” The EMBO Journal 32 (17): 2407–17.

16. Galindo-González, Leonardo, Corinne Mhiri, Michael K. Deyholos, and Marie Angèle Grandbastien. 2017. “LTR-Retrotransposons in Plants: Engines of Evolution.” Gene 626 (May): 14–25.

17. Grant, Charles E., Timothy L. Bailey, and William Stafford Noble. 2011. “FIMO: Scanning for Occurrences of a given Motif.” Bioinformatics 27 (7): 1017–18.

18. Guo, Yuchun, Shaun Mahony, and David K. Gifford. 2012. “High Resolution Genome Wide Binding Event Finding and Motif Discovery Reveals Transcription Factor Spatial Binding Constraints.” PLoS Computational Biology 8 (8): e1002638.

19. Hershberger, R. J., M. I. Benito, K. J. Hardeman, C. Warren, V. L. Chandler, and V. Walbot. 1995. “Characterization of the Major Transcripts Encoded by the Regulatory MuDR Transposable Element of Maize.” Genetics 140 (3): 1087–98.

20. Hosaka, Aoi, Raku Saito, Kazuya Takashima, Taku Sasaki, Yu Fu, Akira Kawabe, Tasuku Ito, et al. 2017. “Evolution of Sequence-Specific Anti-Silencing Systems in Arabidopsis.” Nature Communications 8 (1): 1–10.

21. Huang, Shengfeng, Xin Tao, Shaochun Yuan, Yuhang Zhang, Peiyi Li, Helen A. Beilinson, Ya Zhang, et al. 2016. “Discovery of an Active RAG Transposon Illuminates the Origins of V(D)J Recombination.” Cell 166 (1): 102–14.

22. Ikeda, Yoko, Thierry Pélissier, Pierre Bourguet, Claude Becker, Marie-Noëlle Pouch-Pélissier, Romain Pogorelcnik, Magdalena Weingartner, Detlef Weigel, Jean-Marc Deragon, and Olivier Mathieu. 2017. “Arabidopsis Proteins with a Transposon-Related Domain Act in Gene Silencing.” Nature Communications 8 (May): 15122.

23. Illergård, Kristoffer, David H. Ardell, and Arne Elofsson. 2009. “Structure Is Three to Ten Times More Conserved than Sequence--a Study of Structural Response in Protein Cores.” Proteins 77 (3): 499–508.

24. Ito, Hidetaka, Hervé Herve Gaubert, Etienne Bucher, Marie Mirouze, Isabelle Vaillant, Jerzy Paszkowski, Hervé Herve Gaubert, et al. 2011. “An siRNA Pathway Prevents Transgenerational Retrotransposition in Plants Subjected to Stress.” Nature 472 (7341): 115– 19.

25. Jiang, Ning, Zhirong Bao, Xiaoyu Zhang, Sean R. Eddy, and Susan R. Wessler. 2004. “Pack-MULE Transposable Elements Mediate Gene Evolution in Plants.” Nature 431 (7008): 569–73.

26. Jumper, John, Richard Evans, Alexander Pritzel, Tim Green, Michael Figurnov, Olaf Ronneberger, Kathryn Tunyasuvunakool, et al. 2021. “Highly Accurate Protein Structure Prediction with AlphaFold.” *Nature*, July. 10.1038/s41586-021-03819-2.

27. Kapitonov, Vladimir V., and Jerzy Jurka. 2005. “RAG1 Core and V(D)J Recombination Signal Sequences Were Derived from Transib Transposons.” PLoS Biology 3 (6): e181.

28. Kapitonov, V. V., and J. Jurka. 1999. “Molecular Paleontology of Transposable Elements from Arabidopsis Thaliana.” Genetica 107 (1-3): 27–37.

29. Katoh, Kazutaka, and Daron M. Standley. 2013. “MAFFT Multiple Sequence Alignment Software Version 7: Improvements in Performance and Usability.” Molecular Biology and Evolution 30 (4): 772–80.

30. Kempen, Michel van, Stephanie S. Kim, Charlotte Tumescheit, Milot Mirdita, Jeongjae Lee, Cameron L. M. Gilchrist, Johannes Söding, and Martin Steinegger. 2023. “Fast and Accurate Protein Structure Search with Foldseek.” *Nature Biotechnology*, May. 10.1038/s41587-023-01773-0.

31. Kirchner, J., and S. Sandmeyer. 1993. “Proteolytic Processing of Ty3 Proteins Is Required for Transposition.” Journal of Virology 67 (1): 19–28.

32. Kyte, J., and R. F. Doolittle. 1982. “A Simple Method for Displaying the Hydropathic Character of a Protein.” Journal of Molecular Biology 157 (1): 105–32.

33. Langfelder, Peter, and Steve Horvath. 2008. “WGCNA: An R Package for Weighted Correlation Network Analysis.” BMC Bioinformatics 9 (1): 1–13.

34. Langmead, Ben, and Steven L. Salzberg. 2012. “Fast Gapped-Read Alignment with Bowtie 2.” Nature Methods 9 (4): 357–59.

35. Liao, Yang, Gordon K. Smyth, and Wei Shi. 2014. “featureCounts: An Efficient General Purpose Program for Assigning Sequence Reads to Genomic Features.” Bioinformatics 30 (7): 923–30.

36. Lieber, Michael R., Yunmei Ma, Ulrich Pannicke, and Klaus Schwarz. 2004. “The Mechanism of Vertebrate Nonhomologous DNA End Joining and Its Role in V(D)J Recombination.” DNA Repair 3 (8-9): 817–26.

37. Li, Heng. 2018. “Minimap2: Pairwise Alignment for Nucleotide Sequences.” Bioinformatics 34 (18): 3094–3100.

38. Lin, Rongcheng, Lei Ding, Claudio Casola, Daniel R. Ripoll, Cédric Feschotte, and Haiyang Wang. 2007. “Transposase-Derived Transcription Factors Regulate Light Signaling in Arabidopsis.” Science 318 (5854): 1302–5.

39. Love, Michael I., Wolfgang Huber, and Simon Anders. 2014. “Moderated Estimation of Fold Change and Dispersion for RNA-Seq Data with DESeq2.” Genome Biology 15 (12): 550.

40. Lu, Shennan, Jiyao Wang, Farideh Chitsaz, Myra K. Derbyshire, Renata C. Geer, Noreen R. Gonzales, Marc Gwadz, et al. 2020. “CDD/SPARCLE: The Conserved Domain Database in 2020.” Nucleic Acids Research 48 (D1): D265–68.

41. Masson, Patrick, George Rutherford, Jo Ann Banks, and Nina Fedoroff. 1989. “Essential Large Transcripts of the Maize Spm Transposable Element Are Generated by Alternative Splicing.” Cell 58 (4): 755–65.

42. Material, Supporting Online, Science Web, Highwire Press, New York, and Avenue Nw. 2011. “Evidence for Network Evolution in an Arabidopsis Interactome Map.” Science 333 (6042): 601–7.

43. Merkulov, G. V., K. M. Swiderek, C. B. Brachmann, and J. D. Boeke. 1996. “A Critical Proteolytic Cleavage Site near the C Terminus of the Yeast Retrotransposon Ty1 Gag Protein.” Journal of Virology 70 (8): 5548–56.

44. Mirdita, Milot, Konstantin Schütze, Yoshitaka Moriwaki, Lim Heo, Sergey Ovchinnikov, and Martin Steinegger. 2022. “ColabFold: Making Protein Folding Accessible to All.” Nature Methods 19 (6): 679–82.

45. Moore, S. P., L. A. Rinckel, and D. J. Garfinkel. 1998. “A Ty1 Integrase Nuclear Localization Signal Required for Retrotransposition.” Molecular and Cellular Biology 18 (2): 1105–14.

46. Mukherjee, Krishanu, Luciano Brocchieri, and Thomas R. Bürglin. 2009. “A Comprehensive Classification and Evolutionary Analysis of Plant Homeobox Genes.” Molecular Biology and Evolution 26 (12): 2775–94.

47. Nguyen, Phong Quoc, Christine Conesa, Elise Rabut, Gabriel Bragagnolo, Célia Gouzerh, Carlos Fernández-Tornero, Pascale Lesage, Juan Reguera, and Joël Acker. 2021. “Ty1 Integrase Is Composed of an Active N-Terminal Domain and a Large Disordered C-Terminal Module Dispensable for Its Activity in Vitro.” The Journal of Biological Chemistry 297 (4): 101093.

48. Noé, Laurent, and Gregory Kucherov. 2005. “YASS: Enhancing the Sensitivity of DNA Similarity Search.” Nucleic Acids Research 33 (Web Server issue): W540–43.

49. Oberlin, Stefan, Alexis Sarazin, Clément Chevalier, Olivier Voinnet, and Arturo Marí-Ordóñez. 2017. “A Genome-Wide Transcriptome and Translatome Analysis of Arabidopsis Transposons Identifies a Unique and Conserved Genome Expression Strategy for Ty1/Copia Retroelements.” Genome Research 27 (9): 1549–62.

50. O’Malley, Ronan C., Shao-Shan Carol Huang, Liang Song, Mathew G. Lewsey, Anna Bartlett, Joseph R. Nery, Mary Galli, Andrea Gallavotti, and Joseph R. Ecker. 2016. “Cistrome and Epicistrome Features Shape the Regulatory DNA Landscape.” Cell 165 (5): 1280–92.

51. Panda, Kaushik, and R. Keith Slotkin. 2020. “Long-Read cDNA Sequencing Enables a ‘Gene-Like’ Transcript Annotation of Transposable Elements.” The Plant Cell 32 (9): 2687–98.

52. Price, Morgan N., Paramvir S. Dehal, and Adam P. Arkin. 2010. “FastTree 2--Approximately Maximum-Likelihood Trees for Large Alignments.” PloS One 5 (3): e9490.

53. Quadrana, Leandro, Mathilde Etcheverry, Arthur Gilly, Erwann Caillieux, Mohammed-Amin Madoui, Julie Guy, Amanda Bortolini Silveira, et al. 2019. “Transposition Favors the Generation of Large Effect Mutations That May Facilitate Rapid Adaption.” Nature Communications 10 (1): 3421.

54. Raina, R., M. Schlappi, B. Karunanandaa, A. Elhofy, and N. Fedoroff. 1998. “Concerted Formation of Macromolecular Suppressor-Mutator Transposition Complexes.” Proceedings of the National Academy of Sciences of the United States of America 95 (15): 8526–31.

55. Sasaki, Taku, Kyudo Ro, Erwann Caillieux, Riku Manabe, Grégoire Bohl-Viallefond, Pierre Baduel, Vincent Colot, Tetsuji Kakutani, and Leandro Quadrana. 2022. “Fast Co-Evolution of Anti-Silencing Systems Shapes the Invasiveness of Mu-like DNA Transposons in Eudicots.” The EMBO Journal 41 (8): e110070.

56. Shannon, Paul, Andrew Markiel, Owen Ozier, Nitin S. Baliga, Jonathan T. Wang, Daniel Ramage, Nada Amin, Benno Schwikowski, and Trey Ideker. 2003. “Cytoscape: A Software Environment for Integrated Models of Biomolecular Interaction Networks.” Genome Research 13 (11): 2498–2504.

57. Slotkin, R. Keith, Matthew Vaughn, Filipe Borges, Miloš Tanurdžić, Jörg D. Becker, José A. Feijó, and Robert A. Martienssen. 2009. “Epigenetic Reprogramming and Small RNA Silencing of Transposable Elements in Pollen.” Cell 136 (3): 461–72.

58. Tang, Alison D., Cameron M. Soulette, Marijke J. van Baren, Kevyn Hart, Eva Hrabeta-Robinson, Catherine J. Wu, and Angela N. Brooks. 2020. “Full-Length Transcript Characterization of SF3B1 Mutation in Chronic Lymphocytic Leukemia Reveals Downregulation of Retained Introns.” Nature Communications 11 (1): 1438.

59. Varadi, Mihaly, Stephen Anyango, Mandar Deshpande, Sreenath Nair, Cindy Natassia, Galabina Yordanova, David Yuan, et al. 2022. “AlphaFold Protein Structure Database: Massively Expanding the Structural Coverage of Protein-Sequence Space with High-Accuracy Models.” Nucleic Acids Research 50 (D1): D439–44.

60. West, Alan Mv, Scott C. Rosenberg, Sarah N. Ur, Madison K. Lehmer, Qiaozhen Ye, Götz Hagemann, Iracema Caballero, et al. 2019. “A Conserved Filamentous Assembly Underlies the Structure of the Meiotic Chromosome Axis.” eLife 8 (January). 10.7554/eLife.40372.

61. Wicker, Thomas, Romain Guyot, Nabila Yahiaoui, and Beat Keller. 2003. “CACTA Transposons in Triticeae. A Diverse Family of High-Copy Repetitive Elements.” Plant Physiology 132 (1): 52– 63.

62. Wicker, Thomas, François Sabot, Aurélie Hua-Van, Jeffrey L. Bennetzen, Pierre Capy, Boulos Chalhoub, Andrew Flavell, et al. 2007. “A Unified Classification System for Eukaryotic Transposable Elements.” Nature Reviews. Genetics 8 (12): 973–82.

63. Wlodzimierz, Piotr, Fernando A. Rabanal, Robin Burns, Matthew Naish, Elias Primetis, Alison Scott, Terezie Mandáková, et al. 2023. “Cycles of Satellite and Transposon Evolution in Arabidopsis Centromeres.” Nature 618 (7965): 557–65.

64. Youkharibache, Philippe, Stella Veretnik, Qingliang Li, Kimberly A. Stanek, Cameron Mura, and Philip E. Bourne. 2019. “The Small β-Barrel Domain: A Survey-Based Structural Analysis.” Structure 27 (1): 6–26.

65. Youngren, S. D., J. D. Boeke, N. J. Sanders, and D. J. Garfinkel. 1988. “Functional Organization of the Retrotransposon Ty from Saccharomyces Cerevisiae: Ty Protease Is Required for Transposition.” Molecular and Cellular Biology 8 (4): 1421–31.

66. Yuan, Qianmu, Sheng Chen, Jiahua Rao, Shuangjia Zheng, Huiying Zhao, and Yuedong Yang. 2022. “AlphaFold2-Aware Protein-DNA Binding Site Prediction Using Graph Transformer.” Briefings in Bioinformatics 23 (2). 10.1093/bib/bbab564.

67. Yu, Guangchuang, Li-Gen Wang, Yanyan Han, and Qing-Yu He. 2012. “clusterProfiler: An R Package for Comparing Biological Themes among Gene Clusters.” Omics: A Journal of Integrative Biology 16 (5): 284–87.

68. Zhang, Runxuan, Cristiane P. G. Calixto, Yamile Marquez, Peter Venhuizen, Nikoleta A. Tzioutziou, Wenbin Guo, Mark Spensley, et al. 2017. “A High Quality Arabidopsis Transcriptome for Accurate Transcript-Level Analysis of Alternative Splicing.” Nucleic Acids Research 45 (9): 5061–73.

69. Zhu, Yujuan, Xiaoying Hu, Ying Duan, Shaofang Li, Yu Wang, Amin Ur Rehman, Junna He, et al. 2020. “The Arabidopsis Nodulin Homeobox Factor AtNDX Interacts with AtRING1A/B and Negatively Regulates Abscisic Acid Signaling.” The Plant Cell 32 (3): 703–21.

70. Zuker, Michael. 2003. “Mfold Web Server for Nucleic Acid Folding and Hybridization Prediction.” Nucleic Acids Research 31 (13): 3406–15.

